## Supplemental Figures for "An endosymbiont harvest: Phylogenomic analysis of *Wolbachia* genomes from the Darwin Tree of Life biodiversity genomics project"

#### Figure S1: Sampling locations and incidence of *Wolbachia*

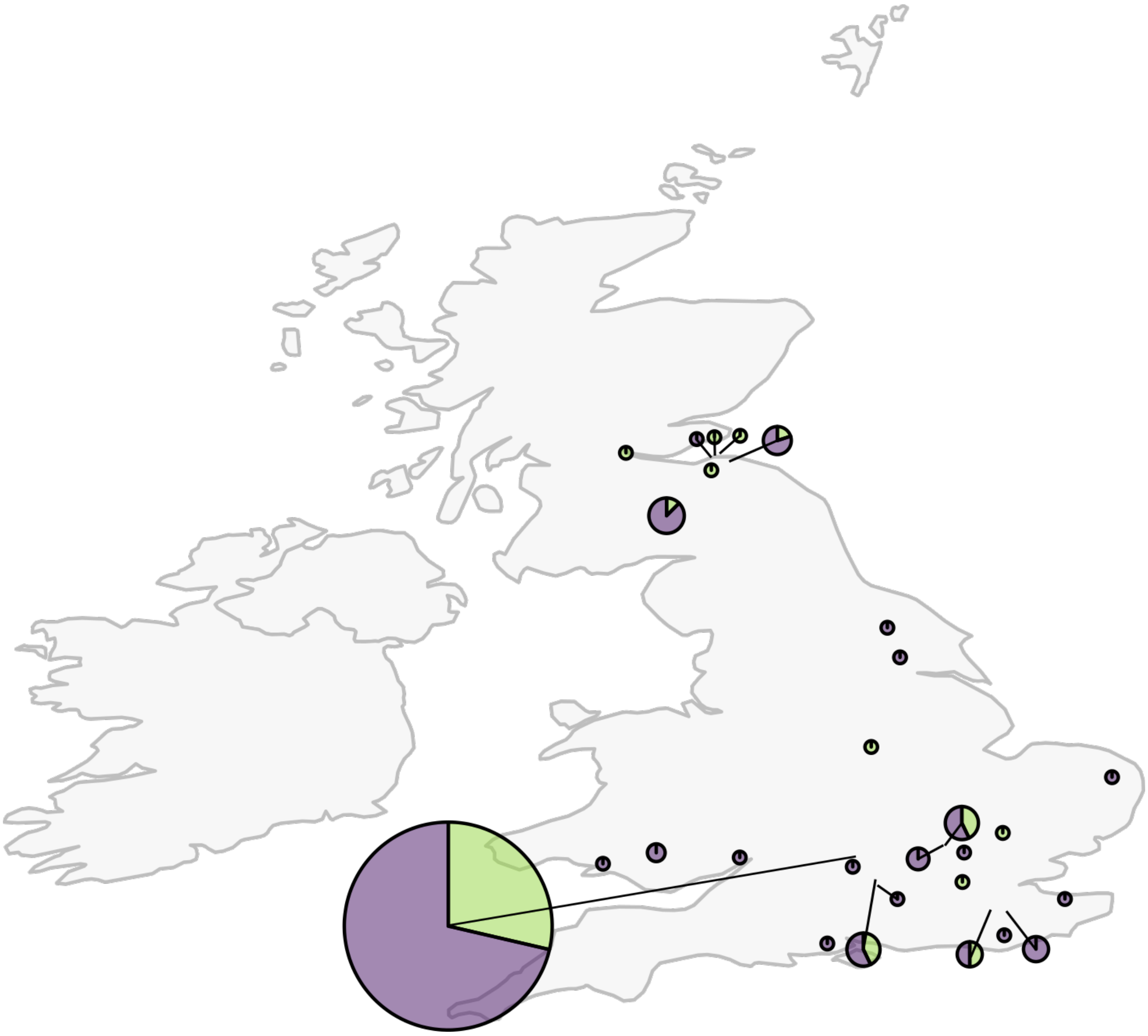

**Figure S1:** Sampling locations and incidence of *Wolbachia* presence (green) and absence (purple) of DToL samples from Britain and Ireland. The size of the pie charts reflect the number of collected samples per location. Most samples came from Wytham Woods Genomic Observatory near Oxford.

#### Figure S2: Contiguity and genome size distribution of *Wolbachia*

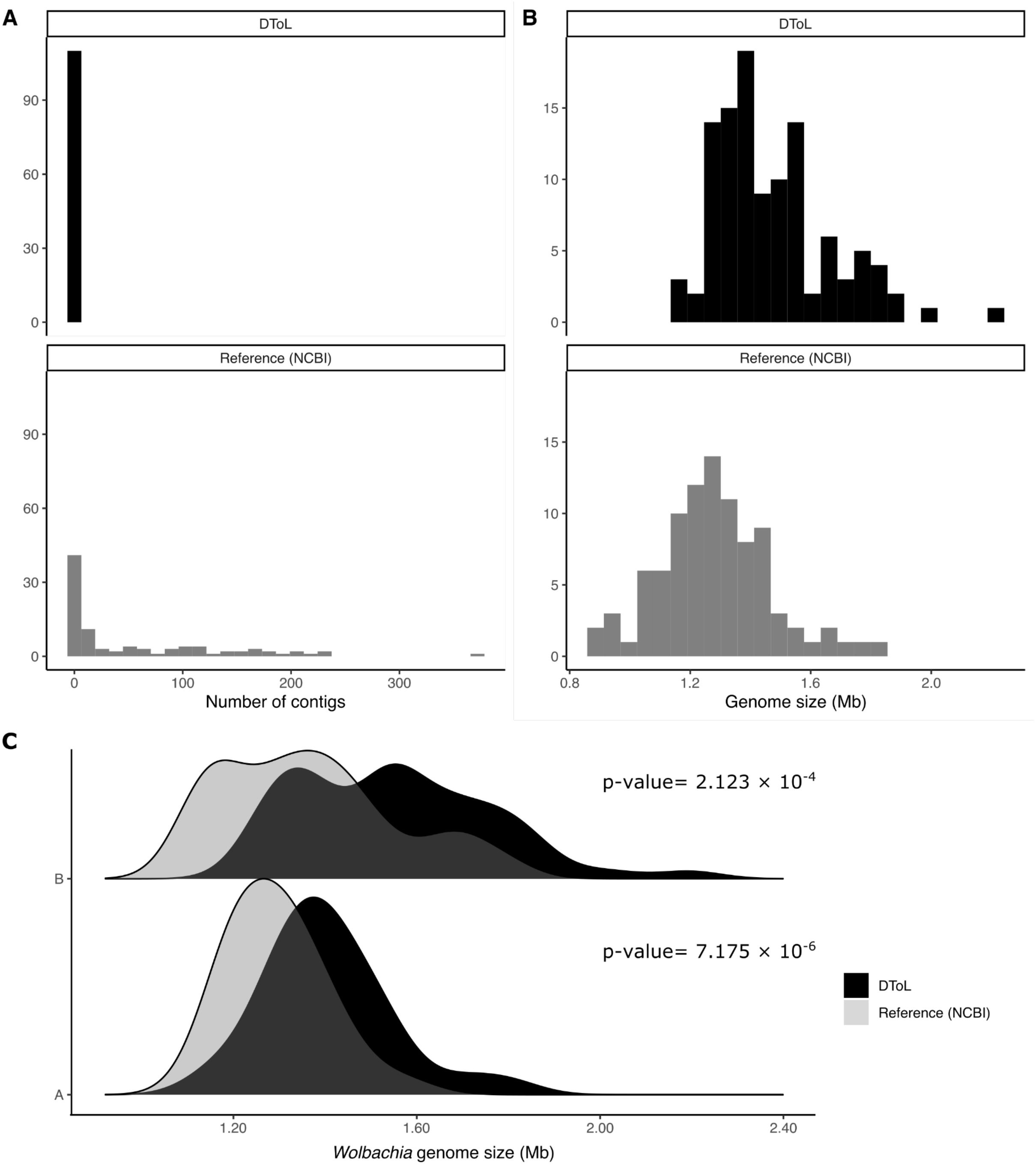


**Figure S2:**
A,B Contiguity and genome size distribution of Wolbachia genomes assembled in this study (black) vs reference genomes from other projects available in NCBI (grey).
C. Genome size distribution of *Wolbachia*. supergroup A (above) and B (below), in this study (black) and reference genomes from other projects available in NCBI (grey).

### Figure S3: Phylogeny of supergroup A and B *Wolbachia*

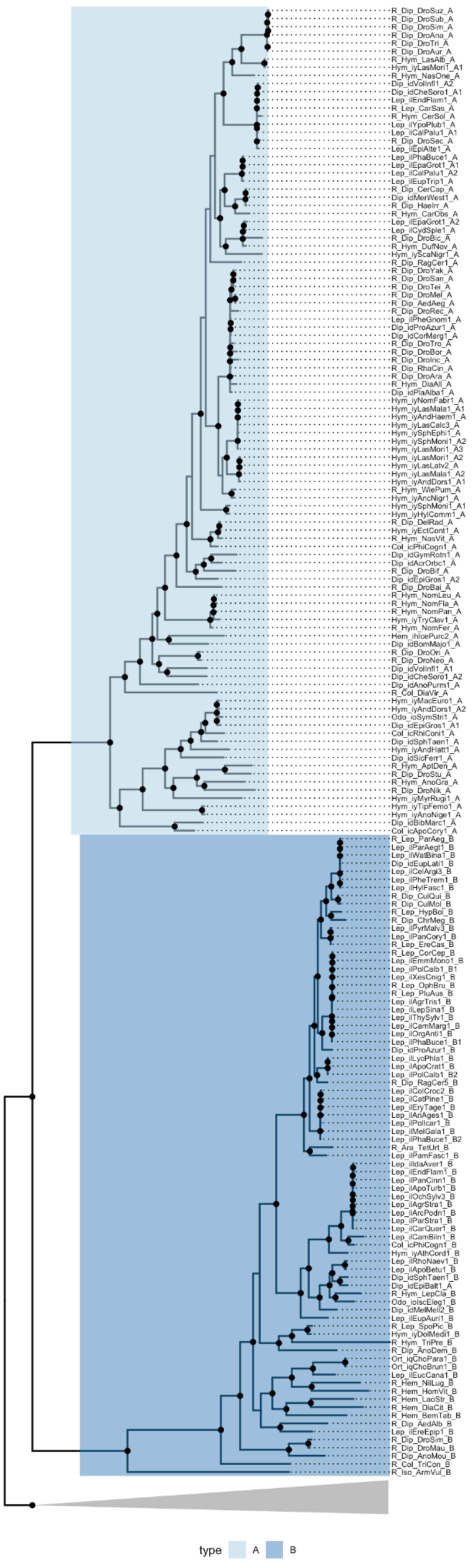


**Figure S3:** Phylogeny of supergroup A and B *Wolbachia*, visualised with the root placed between the A and B supergroups and the remaining supergroups (C,D,E,F, J, S; nodes collapsed as grey wedge), highlighting nodes with bootstrap value higher than 80 with a black label.

#### Figure S4: Average nucleotide identity between Wytham Woods specimens

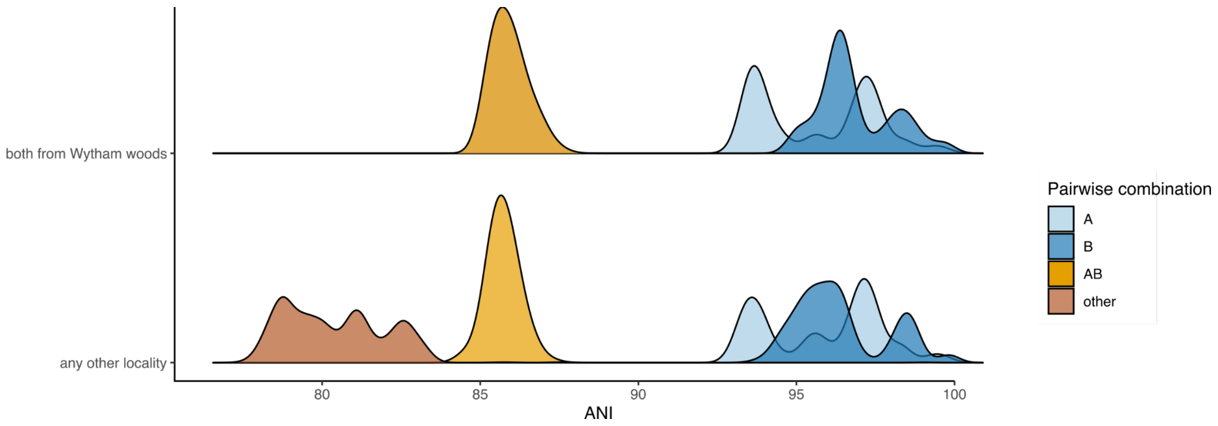


**Figure S4:** Distribution of average nucleotide identity (ANI) between pairs of *Wolbachia* genomes if specimens were both sampled from Wytham Woods (upper panel) or any other locality (lower panel). Distributions are separated by the classification of the two genomes, i.e. both belonging to supergroup A, both belonging to supergroup B, comparisons of A with B, or comparisons between other supergroups.

#### Figure S5: Predicted proteome size in *Wolbachia*

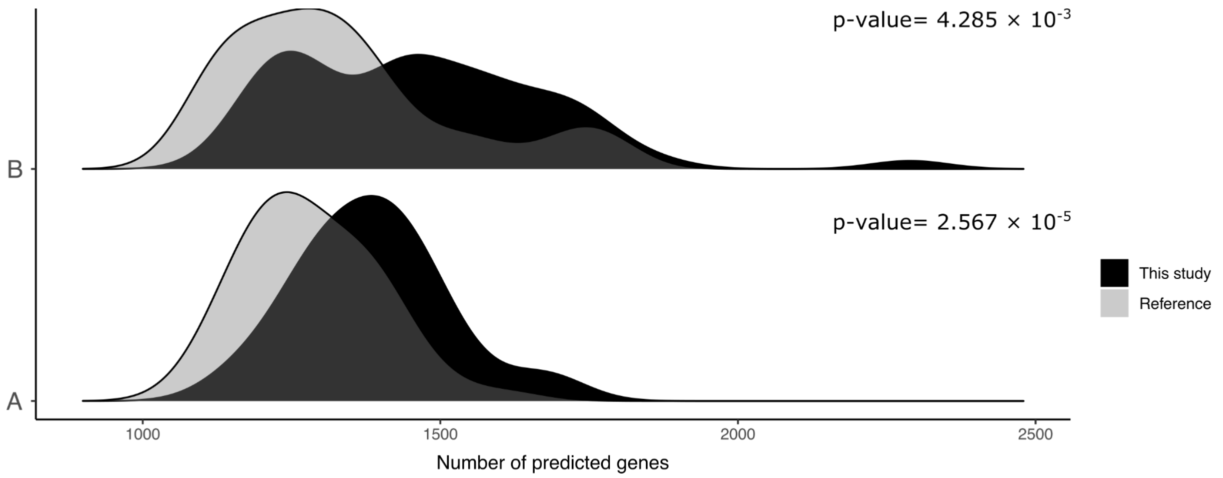


**Figure S5:** Number of predicted protein-coding genes for *Wolbachia* supergroup A (above) and B (below), in this study (black) and reference genomes from other projects available in NCBI (grey).

#### Figure S6: Strain-specific proteins are not generally associated with WO phage.

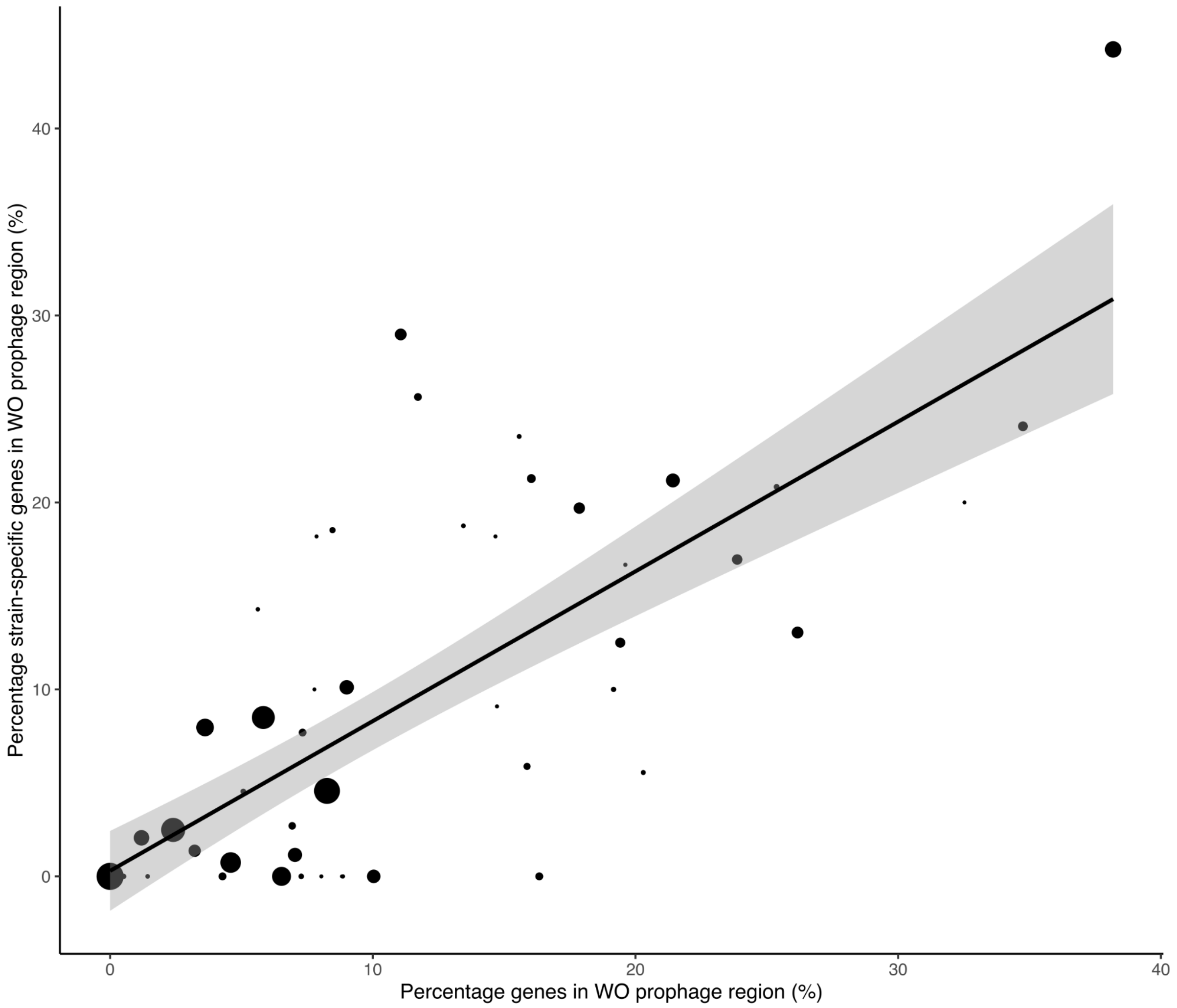


**Figure S6**: Percentage of protein-coding genes present in WO prophage regions versus percentage of strain-specific protein-coding genes in those regions of *Wolbachia* genomes with at least 10 strain-specific genes. Size of points is reflective of the total number of strain-specific genes. Linear regression line with confidence interval is displayed.

#### Figure S7: Comparison of the phylogenies of biotin synthesis clusters and the *Wolbachia* strains that contain them.

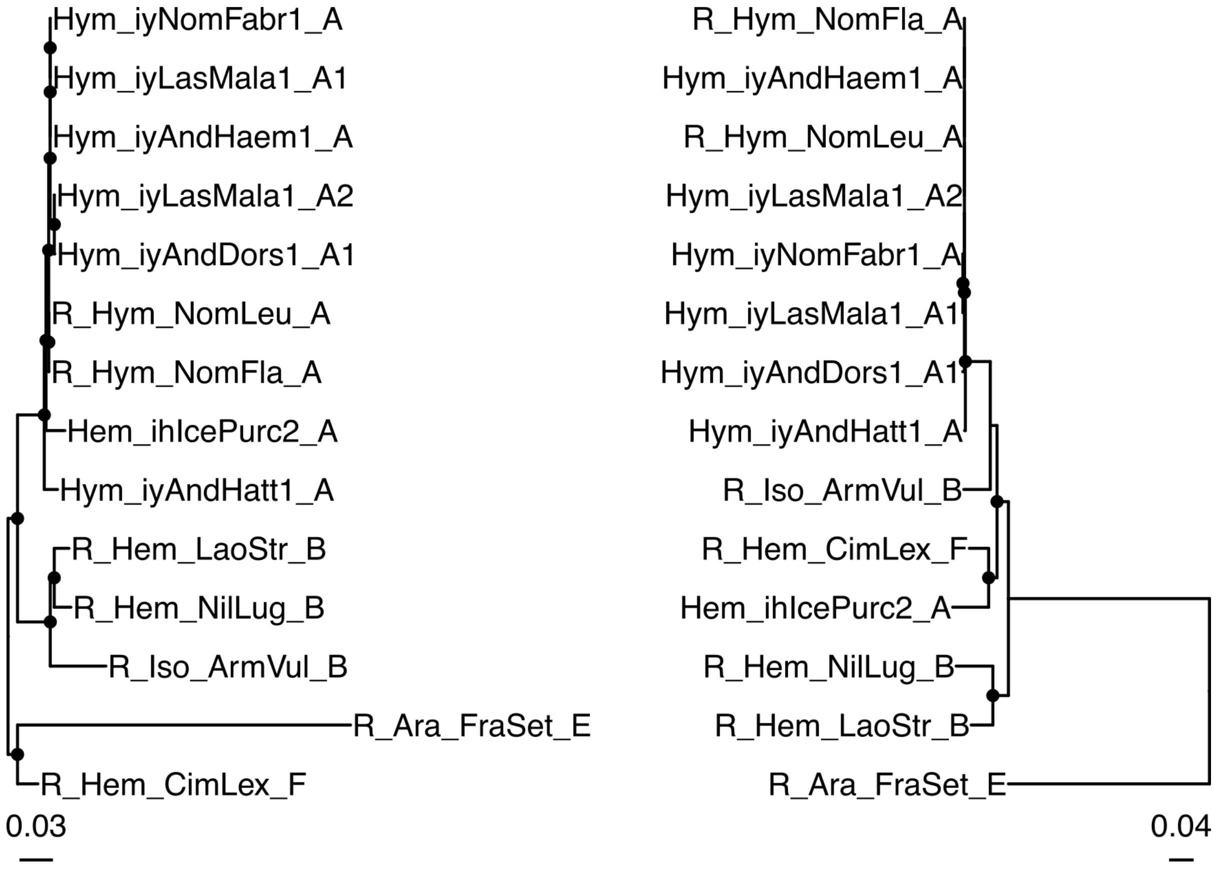


**Figure S7**: Comparison between phylogenies of *Wolbachia* genomes containing the biotin locus, based on tree in Fig2A (left) and a phylogeny inferred from the six nucleotide genes constituting the biotin synthesis operon (BioA-D,BioF, BioH) (right). Internal nodes with bootstrap support higher than 80 are highlighted with black circles.
