## Supplemental Table 1: DToL screened genomes for "An endosymbiont harvest: Phylogenomic analysis of *Wolbachia* genomes from the Darwin Tree of Life biodiversity genomics project"

**Table S1: Overview of all DToL screened genomes**

| Scientific name | Species name | Taxonomic order | Biospecimen ID | Sex |
|---|---|---|---|---|
| *Abrostola tripartita* | The Spectacle | Lepidoptera | SAMEA7520667 | Male |
| *Acleris emargana* | Notch-wing Button | Lepidoptera | SAMEA7746613 | Male |
| *Acleris sparsana* | Ashy button | Lepidoptera | SAMEA8603209 | Male |
| *Acrobasis consociella* | Broad-barred Knot-horn | Lepidoptera | SAMEA7701461 | Unknown |
| *Acrobasis repandana* | Warted Knot-horn | Lepidoptera | SAMEA7701659 | Unknown |
| *Acronicta aceris* | The sycamore | Lepidoptera | SAMEA7701532 | Female |
| *Agonopterix arenella* | Brindled Flat-body | Lepidoptera | SAMEA8603192 | Male |
| *Agonopterix heracliana* | Common flat-body | Lepidoptera | SAMEA7701514 | Unknown |
| *Agonopterix subpropinquella* | Ruddy Flat-body | Lepidoptera | SAMEA7746629 | Male |
| *Agriopis aurantiaria* | Scarce umber | Lepidoptera | SAMEA8603214 | Male |
| *Agriphila geniculea* | Elbow-stripe Grass-veneer | Lepidoptera | SAMEA8603180 | Female |
| *Agriphila straminella* | Straw Grass-veneer | Lepidoptera | SAMEA7701512 | Unknown |
| *Agriphila tristella* | Common grass-veneer | Lepidoptera | SAMEA8603174 | Male |
| *Agrochola circellaris* | The brick | Lepidoptera | SAMEA8603201 | Male |
| *Agrochola macilenta* | Yellow-line quaker | Lepidoptera | SAMEA8603207 | Female |
| *Allophyes oxyacanthae* | Green-brindled crescent | Lepidoptera | SAMEA8603204 | Male |
| *Amphipyra berbera* | Svensson's copper underwing | Lepidoptera | SAMEA7701493 | Male |
| *Amphipyra tragopoginis* | Mouse moth | Lepidoptera | SAMEA7520174 | Male |
| *Anthocharis cardamines* | Orange tip | Lepidoptera | SAMEA7523110 | Female |
| *Apamea monoglypha* | Dark arches | Lepidoptera | SAMEA7701555 | Male |
| *Apatura iris* | Purple emperor | Lepidoptera | SAMEA7523112 | Unknown |
| *Apeira syringaria* | Lilac beauty | Lepidoptera | SAMEA7520685 | Male |
| *Aplocera efformata* | Lesser treble-bar | Lepidoptera | SAMEA8603170 | Female |
| *Aporia crataegi* | Black Veined White | Lepidoptera | SAMEA7523355 | Male |
| *Aporophyla lueneburgensis* | Northern deep-brown dart | Lepidoptera | SAMEA8603194 | Female |
| *Apotomis betuletana* | Birch marble | Lepidoptera | SAMEA7701588 | Male |
| *Apotomis turbidana* | White-shouldered Marble | Lepidoptera | SAMEA7520681 | Unknown |

| | | | | |
|---|---|---|---|---|
| *Archips podanus* | Large Fruit-tree Tortrix | Lepidoptera | SAMEA7701540 | Unknown |
| *Archips xylosteana* | Variegated Golden Tortrix | Lepidoptera | SAMEA7701541 | Unknown |
| *Argyresthia goedartella* | Golden Argent | Lepidoptera | SAMEA7520176 | Unknown |
| *Aricia agestis* | Brown argus | Lepidoptera | SAMEA7523300 | Male |
| *Atethmia centrago* | Centre-barred sallow | Lepidoptera | SAMEA7520177 | Male |
| *Autographa gamma* | Silver Y | Lepidoptera | SAMEA7519848 | Female |
| *Autographa pulchrina* | Beautiful Golden Y | Lepidoptera | SAMEA7520527 | Female |
| *Bembecia ichneumoniformis* | Six-belted clearwing | Lepidoptera | SAMEA7701282 | Male |
| *Biston betularia* | Peppered moth | Lepidoptera | SAMEA7520512 | Male |
| *Blastobasis adustella* | Dingy Dowd | Lepidoptera | SAMEA7520179 | Female |
| *Blastobasis lacticolella* | London Dowd | Lepidoptera | SAMEA7519826 | Male |
| *Boloria selene* | Small pearl-bordered fritillary | Lepidoptera | SAMEA7523131 | Female |
| *Calamotropha paludella* | Bulrush Veneer | Lepidoptera | SAMEA7746625 | Male |
| *Campaea margaritaria* | Light emerald | Lepidoptera | SAMEA7701535 | Male |
| *Camptogramma bilineatum* | Yellow shell | Lepidoptera | SAMEA7701528 | Unknown |
| *Caradrina clavipalpis* | Pale mottled willow | Lepidoptera | SAMEA8603187 | Male |
| *Caradrina kadenii* | Clancy’s Rustic | Lepidoptera | SAMEA8534280 | Female |
| *Carcina quercana* | Long-horned flat-body | Lepidoptera | SAMEA7519850 | Male |
| *Catocala fraxini* | Cliften non-pareil | Lepidoptera | SAMEA8603175 | Male |
| *Catoptria pinella* | Pearl Grass-veneer | Lepidoptera | SAMEA7701506 | Unknown |
| *Celastrina argiolus* | Holly brown | Lepidoptera | SAMEA7523268 | Male |
| *Chloroclysta siterata* | Red-green carpet | Lepidoptera | SAMEA8603199 | Male |
| *Chloroclystis v-ata* | V-pug | Lepidoptera | SAMEA7701460 | Unknown |
| *Chrysoteuchia culmella* | Garden Grass-veneer | Lepidoptera | SAMEA7701502 | Male |
| *Clostera curtula* | Chocolate tip | Lepidoptera | SAMEA7520526 | Female |
| *Colias croceus* | Clouded Yellow | Lepidoptera | SAMEA7523360 | Female |
| *Colostygia pectinataria* | Green carpet | Lepidoptera | SAMEA7520182 | Unknown |
| *Cosmia trapezina* | The dun-bar | Lepidoptera | SAMEA7519851 | Male |
| *Craniophora ligustri* | The coronet | Lepidoptera | SAMEA7519852 | Female |
| *Crocallis elinguaria* | Scalloped oak | Lepidoptera | SAMEA7701527 | Female |

| | | | | |
|---|---|---|---|---|
| *Cupido minimus* | Small blue | Lepidoptera | SAMEA7523306 | Male |
| *Cyaniris semiargus* | Mazarine blue | Lepidoptera | SAMEA7523311 | Male |
| *Cydia fagiglandana* | Large Beech Piercer | Lepidoptera | SAMEA7701453 | Unknown |
| *Cydia splendana* | Marbled Piercer | Lepidoptera | SAMEA7701547 | Female |
| *Deilephila porcellus* | Small elephant Hawk-moth | Lepidoptera | SAMEA7520522 | Male |
| *Diachrysia chrysitis* | Burnished brass moth | Lepidoptera | SAMEA8603181 | Male |
| *Diarsia rubi* | Small square-spot | Lepidoptera | SAMEA8603186 | Female |
| *Ditula angustiorana* | Red-barred Tortrix | Lepidoptera | SAMEA7701319 | Unknown |
| *Dryobotodes eremita* | Brindled green | Lepidoptera | SAMEA8603190 | Female |
| *Ecliptopera silaceata* | Small phoenix | Lepidoptera | SAMEA7701534 | Male |
| *Ectropis crepuscularia* | The engrailed | Lepidoptera | SAMEA7701526 | Unknown |
| *Eilema depressum* | Buff footman | Lepidoptera | SAMEA7746611 | Male |
| *Eilema sororcula* | Orange Footman | Lepidoptera | SAMEA7631555 | Male |
| *Emmelina monodactyla* | Common plume | Lepidoptera | SAMEA8603203 | Female |
| *Endotricha flammealis* | Rosy tabby | Lepidoptera | SAMEA7519855 | Female |
| *Ennomos fuscantarius* | Dusky thorn | Lepidoptera | SAMEA7520185 | Male |
| *Ennomos quercinarius* | August thorn | Lepidoptera | SAMEA7701560 | Male |
| *Epagoge grotiana* | Brown-barred Tortrix | Lepidoptera | SAMEA7701542 | Unknown |
| *Epinotia brunnichana* | Large Birch Bell | Lepidoptera | SAMEA7701529 | Unknown |
| *Epirrhoe alternata* | common carpet | Lepidoptera | SAMEA7701550 | Unknown |
| *Erannis defoliaria* | Mottled umber | Lepidoptera | SAMEA7520367 | Male |
| *Erebia aethiops* | Scotch argus | Lepidoptera | SAMEA7523289 | Female |
| *Erebia ligea* | Mountain ringlet | Lepidoptera | SAMEA7523313 | Male |
| *Erynnis tages* | Dingy skipper | Lepidoptera | SAMEA7523299 | Male |
| *Eucosma campoliliana* | Marbled Bell | Lepidoptera | SAMEA7701295 | Unknown |
| *Eucosma cana* | Hoary Bell | Lepidoptera | SAMEA7701552 | Unknown |
| *Eulithis prunata* | The Pheonix | Lepidoptera | SAMEA7701309 | Male |
| *Euphydryas aurinia* | Marsh Fritillary | Lepidoptera | SAMEA7523466 | Unknown |
| *Eupithecia centaureata* | Lime-speck pug | Lepidoptera | SAMEA7520186 | Male |
| *Eupithecia tripunctaria* | White-spotted pug | Lepidoptera | SAMEA7701546 | Unknown |

| | | | | |
|---|---|---|---|---|
| *Euplexia lucipara* | Small angle shades | Lepidoptera | SAMEA7701470 | Male |
| *Euproctis similis* | Yellow-tail | Lepidoptera | SAMEA7519909 | Male |
| *Eupsilia transversa* | satellite moth | Lepidoptera | SAMEA8563699 | Female |
| *Fabriciana adippe* | High brown fritillary | Lepidoptera | SAMEA7523308 | Female |
| *Furcula furcula* | Sallow kitten | Lepidoptera | SAMEA7746637 | Male |
| *Glaucopsyche alexis* | Green-underside blue | Lepidoptera | SAMEA7524616 | Male |
| *Griposia aprilina* | Merveille du jour | Lepidoptera | SAMEA8603200 | Female |
| *Gymnoscelis rufifasciata* | Double-striped pug | Lepidoptera | SAMEA7519910 | Female |
| *Habrosyne pyritoides* | Buff arches | Lepidoptera | SAMEA7701298 | Male |
| *Hamearis lucina* | Duke of Burgundy | Lepidoptera | SAMEA7523117 | Unknown |
| *Hecatera dysodea* | Small ranunculus | Lepidoptera | SAMEA7521514 | Female |
| *Hedya salicella* | White-backed Marble | Lepidoptera | SAMEA7520688 | Male |
| *Hemaris fuciformis* | broad-bordered bee hawk-moth | Lepidoptera | SAMEA5248724 | Unknown |
| *Hesperia comma* | Silver-spotted skipper | Lepidoptera | SAMEA7523119 | Female |
| *Hydraecia micacea* | Rosy rustic | Lepidoptera | SAMEA8603188 | Female |
| *Hydriomena furcata* | July highflyer | Lepidoptera | SAMEA7701301 | Male |
| *Hylaea fasciaria* | Barred red | Lepidoptera | SAMEA7520684 | Male |
| *Hypena proboscidalis* | The snout | Lepidoptera | SAMEA7520188 | Female |
| *Idaea aversata* | Riband Wave | Lepidoptera | SAMEA7519834 | Male |
| *Laothoe populi* | Poplar Hawk-moth | Lepidoptera | SAMEA7520519 | Female |
| *Lasiommata megera* | Wall brown | Lepidoptera | SAMEA7523153 | Female |
| *Laspeyria flexula* | Beautiful hook-tip | Lepidoptera | SAMEA7519836 | Male |
| *Leptidea sinapis* | Wood White | Lepidoptera | SAMEA7523467 | Male |
| *Limenitis camilla* | White admiral | Lepidoptera | SAMEA7523310 | Female |
| *Lobophora halterata* | The Seraphim | Lepidoptera | SAMEA7520514 | Female |
| *Luperina testacea* | Flounced Rustic | Lepidoptera | SAMEA8534287 | Male |
| *Lycaena phlaeas* | Small copper | Lepidoptera | SAMEA7523293 | Male |
| *Lymantria monacha* | Black arches | Lepidoptera | SAMEA7519912 | Male |
| *Lysandra bellargus* | Adonis Blue | Lepidoptera | SAMEA7523471 | Female |
| *Lysandra coridon* | Chalkhill blue | Lepidoptera | SAMEA7523305 | Male |

| | | | | |
|---|---|---|---|---|
| *Macaria notata* | Peacock moth | Lepidoptera | SAMEA7746623 | Male |
| *Mamestra brassicae* | Cabbage moth | Lepidoptera | SAMEA7524129 | Male |
| *Maniola jurtina* | Meadow brown | Lepidoptera | SAMEA7523158 | Female |
| *Marasmarcha lunaedactyla* | Crescent plume | Lepidoptera | SAMEA7701293 | Female |
| *Meganola albula* | Kent black arches | Lepidoptera | SAMEA7701294 | Male |
| *Melanargia galathea* | Marbled white | Lepidoptera | SAMEA7523296 | Female |
| *Melitaea cinxia* | Glanville Fritillary | Lepidoptera | SAMEA7523475 | Male |
| *Mellicta athalia* | Heath fritillary | Lepidoptera | SAMEA7523312 | Female |
| *Mesoligia furuncula* | Cloaked minor | Lepidoptera | SAMEA7701289 | Female |
| *Mimas tiliae* | Lime Hawk-moth | Lepidoptera | SAMEA7520521 | Male |
| *Mythimna albipuncta* | White-point | Lepidoptera | SAMEA8603191 | Male |
| *Mythimna ferrago* | The clay | Lepidoptera | SAMEA7701536 | Female |
| *Mythimna impura* | Smoky wainscot | Lepidoptera | SAMEA7519913 | Female |
| *Neocochylis molliculana* | Ox-tongue Conch | Lepidoptera | SAMEA7746615 | Unknown |
| *Noctua comes* | Lesser yellow underwing | Lepidoptera | SAMEA7701458 | Unknown |
| *Noctua fimbriata* | Broad-bordered yellow underwing | Lepidoptera | SAMEA7519914 | Female |
| *Noctua janthe* | Lesser broad-boardered yellow underwing | Lepidoptera | SAMEA7701537 | Male |
| *Noctua pronuba* | Large Yellow Underwing | Lepidoptera | SAMEA7519837 | Female |
| *Notocelia uddmanniana* | Bramble shoot moth | Lepidoptera | SAMEA7519916 | Male |
| *Notodonta dromedarius* | Iron prominent | Lepidoptera | SAMEA7520190 | Male |
| *Notodonta ziczac* | Pebble prominent | Lepidoptera | SAMEA7746619 | Male |
| *Nymphalis c-album* | Comma | Lepidoptera | SAMEA7523165 | Female |
| *Nymphalis io* | Peacock | Lepidoptera | SAMEA7523149 | Male |
| *Nymphalis polychloros* | Large Tortoiseshell | Lepidoptera | SAMEA7523477 | Female |
| *Nymphalis urticae* | Small tortoiseshell | Lepidoptera | SAMEA7523286 | Female |
| *Ochlodes sylvanus* | Large skipper | Lepidoptera | SAMEA7523138 | Female |
| *Ochropleura plecta* | Flame shoulder | Lepidoptera | SAMEA7520524 | Female |
| *Omphaloscelis lunosa* | Lunar underwing | Lepidoptera | SAMEA8603195 | Female |
| *Operophtera brumata* | Winter moth | Lepidoptera | SAMEA8563695 | Male |
| *Opisthograptis luteolata* | Brimstone moth | Lepidoptera | SAMEA7519838 | Male |

| | | | | |
|---|---|---|---|---|
| *Orgyia antiqua* | Rusty tussock moth | Lepidoptera | SAMEA7524390 | Male |
| *Ostrinia nubilalis* | European corn borer | Lepidoptera | SAMEA7701321 | Unknown |
| *Pammene fasciana* | Acorn Piercer | Lepidoptera | SAMEA7701530 | Male |
| *Pandemis cinnamomeana* | White-faced tortrix | Lepidoptera | SAMEA8603177 | Male |
| *Pandemis corylana* | Chequered Fruit-tree Tortrix | Lepidoptera | SAMEA7701543 | Unknown |
| *Papilio machaon* | common yellow swallowtail | Lepidoptera | SAMEA7523121 | Female |
| *Parapoynx stratiotata* | Ringed china-mark | Lepidoptera | SAMEA7519920 | Male |
| *Pararge aegeria* | Speckled wood | Lepidoptera | SAMEA7532732 | Female |
| *Peribatodes rhomboidaria* | Willow beauty | Lepidoptera | SAMEA7701524 | Male |
| *Perizoma alchemillatum* | Small rivulet | Lepidoptera | SAMEA7701545 | Unknown |
| *Perizoma flavofasciatum* | Sandy carpet | Lepidoptera | SAMEA7701445 | Unknown |
| *Phalera bucephala* | Buff-tip | Lepidoptera | SAMEA7519921 | Female |
| *Pheosia gnoma* | Lesser swallow prominent | Lepidoptera | SAMEA7520513 | Male |
| *Pheosia tremula* | Swallow prominent | Lepidoptera | SAMEA7520523 | Male |
| *Philereme vetulata* | Brown scallop | Lepidoptera | SAMEA7701300 | Female |
| *Phlogophora meticulosa* | Angle shades | Lepidoptera | SAMEA7520192 | Female |
| *Photedes minima* | Small dotted buff | Lepidoptera | SAMEA7701533 | Unknown |
| *Phragmatobia fuliginosa* | Ruby tiger moth | Lepidoptera | SAMEA7701498 | Male |
| *Pieris brassicae* | Large white | Lepidoptera | SAMEA7532735 | Female |
| *Pieris napi* | Green veined white | Lepidoptera | SAMEA7523140 | Male |
| *Pieris rapae* | Small white | Lepidoptera | SAMEA7523164 | Female |
| *Plebejus argus* | Silver-studded blue | Lepidoptera | SAMEA7523294 | Male |
| *Plutella xylostella* | Diamondback moth | Lepidoptera | SAMEA7520369 | Male |
| *Polyommatus icarus* | Common blue | Lepidoptera | SAMEA7523143 | Male |
| *Psoricoptera gibbosella* | Humped Groundling | Lepidoptera | SAMEA7746612 | Unknown |
| *Ptilodon capucinus* | Coxcomb prominent | Lepidoptera | SAMEA7746620 | Male |
| *Ptycholomoides aeriferana* | Yellow Larch Tortrix | Lepidoptera | SAMEA7701538 | Unknown |
| *Pyrgus malvae* | Grizzled skipper | Lepidoptera | SAMEA7523277 | Male |
| *Rhopobota naevana* | Holly tortrix | Lepidoptera | SAMEA7701517 | Unknown |
| *Schrankia costaestrigalis* | Pinion-streaked snout | Lepidoptera | SAMEA7520193 | Male |

| | | | | |
|---|---|---|---|---|
| *Scotopteryx chenopodiata* | Shaded broad-bar | Lepidoptera | SAMEA7701561 | Unknown |
| *Selenia dentaria* | Early thorn | Lepidoptera | SAMEA7701559 | Male |
| *Sesia apiformis* | Hornet moth | Lepidoptera | SAMEA7701281 | Male |
| *Sphinx pinastri* | Pine hawkmoth | Lepidoptera | SAMEA7701449 | Unknown |
| *Spilarctia lutea* | Buff ermine | Lepidoptera | SAMEA7631557 | Female |
| *Spilosoma lubricipeda* | White ermine | Lepidoptera | SAMEA7520525 | Male |
| *Synanthedon vespiformis* | Yellow-legged clearwing | Lepidoptera | SAMEA7701494 | Male |
| *Thumatha senex* | Round-winged muslin | Lepidoptera | SAMEA7701482 | Unknown |
| *Thyatira batis* | Peach blossom | Lepidoptera | SAMEA7519923 | Male |
| *Thymelicus lineola* | Essex skipper | Lepidoptera | SAMEA7523301 | Female |
| *Thymelicus sylvestris* | Small skipper | Lepidoptera | SAMEA7523279 | Male |
| *Tinea semifulvella* | Fulvous Clothes Moth | Lepidoptera | SAMEA7520371 | Male |
| *Tinea trinotella* | Bird's-nest moth | Lepidoptera | SAMEA7519924 | Male |
| *Udea prunalis* | Dusky Pearl | Lepidoptera | SAMEA7701501 | Unknown |
| *Vanessa atalanta* | Red admiral | Lepidoptera | SAMEA7523145 | Female |
| *Vanessa cardui* | Painted lady | Lepidoptera | SAMEA7523147 | Female |
| *Watsonalla binaria* | Oak hook-tip | Lepidoptera | SAMEA7746618 | Female |
| *Xanthorhoe fluctuata* | Garden carpet | Lepidoptera | SAMEA7701525 | Unknown |
| *Xestia c-nigrum* | Setaceous Hebrew character | Lepidoptera | SAMEA8239458 | Male |
| *Xestia xanthographa* | Square-spot rustic | Lepidoptera | SAMEA7520195 | Female |
| *Yponomeuta plumbellus* | Black-tipped Ermine | Lepidoptera | SAMEA7746626 | Unknown |
| *Yponomeuta sedellus* | Grey Ermine | Lepidoptera | SAMEA7746622 | Male |
| *Ypsolopha scabrella* | Wainscot Smudge | Lepidoptera | SAMEA7701504 | Male |
| *Ypsolopha sequella* | Pied smudge | Lepidoptera | SAMEA7519929 | Male |
| *Zeiraphera isertana* | Cock's-head Bell | Lepidoptera | SAMEA7701465 | Unknown |
| *Zeuzera pyrina* | Leopard moth | Lepidoptera | SAMEA7701286 | Male |
| *Zygaena filipendulae* | 6-spot burnet | Lepidoptera | SAMEA7519846 | Female |
| *Apoderus coryli* | Hazel leaf-roller | Coleoptera | SAMEA7520690 | Male |
| *Cantharis rustica* | | Coleoptera | SAMEA7524272 | Male |
| *Podabrus alpinus* | | Coleoptera | SAMEA7520644 | Female |

| | | | | |
|---|---|---|---|---|
| *Rhagonycha fulva* | Common red soldier beetle | Coleoptera | SAMEA7520319 | Female |
| *Dromius quadrimaculatus* | | Coleoptera | SAMEA7520206 | Unknown |
| *Nebria brevicollis* | | Coleoptera | SAMEA7520209 | Female |
| *Nebria salina* | | Coleoptera | SAMEA7524273 | Female |
| *Ophonus ardosiacus* | | Coleoptera | SAMEA7746467 | Female |
| *Pterostichus madidus* | Black clock beetle | Coleoptera | SAMEA7520318 | Male |
| *Adalia bipunctata* | two-spotted ladybird beetle | Coleoptera | SAMEA9089055 | Male |
| *Chilocorus renipustulatus* | kidney-spot ladybird beetle | Coleoptera | SAMEA7520200 | Unknown |
| *Coccinella septempunctata* | Seven-spotted ladybird | Coleoptera | SAMEA7520205 | Female |
| *Harmonia axyridis* | Harlequin ladybird | Coleoptera | SAMEA7520208 | Female |
| *Hippodamia variegata* | Adonis ladybird | Coleoptera | SAMEA7849270 | Unknown |
| *Propylea quattuordecimpunctata* | 14-spot ladybird | Coleoptera | SAMEA7520316 | Unknown |
| *Rhinocyllus conicus* | | Coleoptera | SAMEA7701490 | Unknown |
| *Agrypnus murinus* | | Coleoptera | SAMEA7701277 | Male |
| *Melolontha melolontha* | cockchafer | Coleoptera | SAMEA7524378 | Male |
| *Malachius bipustulatus* | Common malachite beetle | Coleoptera | SAMEA7520537 | Female |
| *Oedemera lurida* | | Coleoptera | SAMEA7520207 | Unknown |
| *Pyrochroa serraticornis* | Red-headed cardinal beetle | Coleoptera | SAMEA7524259 | Male |
| *Phosphuga atrata* | Black snail beetle | Coleoptera | SAMEA7520321 | Unknown |
| *Ocypus olens* | Devil's coach horse | Coleoptera | SAMEA7520211 | Female |
| *Philonthus cognatus* | | Coleoptera | SAMEA8603235 | Male |
| *Acrocera orbiculus* | Top-horned Hunchback | Diptera | SAMEA7701562 | Unknown |
| *Delia platura* | Bean seed fly | Diptera | SAMEA7746747 | Unknown |
| *Machimus atricapillus* | | Diptera | SAMEA7849389 | Male |
| *Bombylius discolor* | dotted bee fly | Diptera | SAMEA7524252 | Female |
| *Bombylius major* | dark edged bee fly | Diptera | SAMEA7524251 | Male |
| *Bellardia pandia* | Bisetose Emerald-bottle | Diptera | SAMEA7746779 | Female |
| *Lucilia richardsi* | | Diptera | SAMEA8603134 | Unknown |
| *Pollenia angustigena* | Narrow-cheeked Clusterfly | Diptera | SAMEA7746597 | Female |
| *Protocalliphora azurea* | Bird blowfly | Diptera | SAMEA7746778 | Male |

| | | | | |
|---|---|---|---|---|
| *Clusia tigrina* | | Diptera | SAMEA7701567 | Male |
| *Physocephala rufipes* | Waisted Beegrabber | Diptera | SAMEA7746469 | Unknown |
| *Sicus ferrugineus* | Ferruginous Bee-grabber | Diptera | SAMEA7520692 | Male |
| *Thecophora atra* | | Diptera | SAMEA7849382 | Male |
| *Tipula paludosa* | European crane fly | Diptera | SAMEA7520335 | Female |
| *Empis livida* | | Diptera | SAMEA8603147 | Unknown |
| *Callomyia amoena* | | Diptera | SAMEA9066034 | Female |
| *Stomorhina lunata* | Locust Blowfly | Diptera | SAMEA7849406 | Female |
| *Tachina fera* | | Diptera | SAMEA7520333 | Female |
| *Sarcophaga caerulescens* | | Diptera | SAMEA7746589 | Male |
| *Sarcophaga crassimargo* | | Diptera | SAMEA7746602 | Male |
| *Sarcophaga rosellei* | | Diptera | SAMEA7746603 | Male |
| *Sarcophaga variegata* | | Diptera | SAMEA8603132 | Male |
| *Scathophaga stercoraria* | yellow dung fly | Diptera | SAMEA7520161 | Male |
| *Coremacera marginata* | | Diptera | SAMEA7521524 | Female |
| *Baccha elongata* | Gossamer Hoverfly | Diptera | SAMEA7520030 | Female |
| *Cheilosia pagana* | Parsley Cheilosia | Diptera | SAMEA7746768 | Female |
| *Cheilosia soror* | Red-horned Truffle Cheilosia | Diptera | SAMEA7520031 | Female |
| *Cheilosia vulpina* | Large Burdock Cheilosia | Diptera | SAMEA7746587 | Female |
| *Chrysotoxum bicinctum* | Two-banded wasp hoverfly | Diptera | SAMEA7520032 | Female |
| *Chrysotoxum verralli* | Verrall's wasp hoverfly | Diptera | SAMEA7520033 | Female |
| *Criorhina berberina* | Dimorphic Bear Hoverfly | Diptera | SAMEA7701563 | Female |
| *Epistrophe grossulariae* | Broad-banded Epistrophe | Diptera | SAMEA8603153 | Female |
| *Episyrphus balteatus* | Marmalade hoverfly | Diptera | SAMEA7520035 | Unknown |
| *Eristalinus sepulchralis* | Small Spotty-eyed Dronefly | Diptera | SAMEA7746477 | Female |
| *Eristalis arbustorum* | Plane-faced dronefly | Diptera | SAMEA7520036 | Female |
| *Eristalis horticola* | | Diptera | SAMEA7702268 | Female |
| *Eristalis pertinax* | Tapered Dronefly | Diptera | SAMEA7520039 | Male |
| *Eristalis tenax* | Common Dronefly | Diptera | SAMEA7520042 | Female |
| *Eupeodes corollae* | | Diptera | SAMEA7524255 | Female |

| | | | | |
|---|---|---|---|---|
| *Eupeodes latifasciatus* | Meadow Field Syrph | Diptera | SAMEA7746776 | Female |
| *Leucozona laternaria* | Dark-saddled Leucozona | Diptera | SAMEA8603164 | Female |
| *Melanostoma mellinum* | Dumpy grass hoverfly | Diptera | SAMEA7520051 | Male |
| *Melanostoma scalare* | Slender grass hoverfly | Diptera | SAMEA7520053 | Male |
| *Myathropa florea* | Batman hoverfly | Diptera | SAMEA7520156 | Male |
| *Platycheirus albimanus* | White-footed hoverfly | Diptera | SAMEA7520157 | Female |
| *Rhingia campestris* | Common Snout-hoverfly | Diptera | SAMEA7520159 | Male |
| *Scaeva pyrastri* | Pied hoverfly | Diptera | SAMEA7520160 | Female |
| *Sphaerophoria taeniata* | | Diptera | SAMEA7746606 | Male |
| *Volucella inanis* | Lesser hornet hoverfly | Diptera | SAMEA7520171 | Female |
| *Volucella inflata* | Cossus Hoverfly | Diptera | SAMEA7701275 | Male |
| *Xanthogramma pedissequum* | Superb ant-hill hoverfly | Diptera | SAMEA7520951 | Male |
| *Xylota sylvarum* | Golden-tailed hoverfly | Diptera | SAMEA7520173 | Male |
| *Cistogaster globosa* | | Diptera | SAMEA7746478 | Male |
| *Gymnosoma rotundatum* | | Diptera | SAMEA7849381 | Male |
| *Nowickia ferox* | | Diptera | SAMEA7746479 | Female |
| *Thecocarcelia acutangulata* | | Diptera | SAMEA7746598 | Female |
| *Anomoia purmunda* | Hawthorn fruitfly | Diptera | SAMEA7520325 | Female |
| *Merzomyia westermanni* | | Diptera | SAMEA7746463 | Unknown |
| *Terellia serratulae* | | Diptera | SAMEA7520334 | Female |
| *Bibio marci* | St Mark's Fly | Diptera | SAMEA7524263 | Male |
| *Nephrotoma flavescens* | Tiger Cranefly | Diptera | SAMEA7520954 | Male |
| *Cloeon dipterum* | Pond Olive | Ephemeroptera | SAMEA7520803 | Unknown |
| *Ecdyonurus torrentis* | Large brook dun | Ephemeroptera | SAMEA7520824 | Unknown |
| *Rhithrogena germanica* | | Ephemeroptera | SAMEA9065858 | Unknown |
| *Acanthosoma haemorrhoidale* | Hawthorn shieldbug | Hemiptera | SAMEA8563710 | Male |
| *Gonocerus acuteangulatus* | Box Bug | Hemiptera | SAMEA7524254 | Unknown |
| *Pantilius tunicatus* | | Hemiptera | SAMEA7520359 | Unknown |
| *Himacerus mirmicoides* | Ant damselbug | Hemiptera | SAMEA7520349 | Unknown |
| *Notonecta glauca* | Backswimmer / Water Boatman | Hemiptera | SAMEA7520812 | Unknown |

| | | | | |
|---|---|---|---|---|
| *Icerya purchasi* | cottony cushion scale | Hemiptera | SAMEA7523480 | Female |
| *Aelia acuminata* | Bishop's mitre shieldbug | Hemiptera | SAMEA7520338 | Male |
| *Eurydema oleracea* | Brassica shieldbug | Hemiptera | SAMEA7701485 | Unknown |
| *Planococcus citri* | citrus mealybug | Hemiptera | SAMEA7523510 | Female |
| *Andrena dorsata* | Short-fringed Mining Bee | Hymenoptera | SAMEA7746464 | Female |
| *Andrena haemorrhoa* | Red-tailed mining bee | Hymenoptera | SAMEA7520535 | Female |
| *Andrena hattorfiana* | Large scabious mining bee | Hymenoptera | SAMEA7746468 | Female |
| *Bombus campestris* | Field cuckoo-bee | Hymenoptera | SAMEA7520482 | Male |
| *Bombus hortorum* | Garden bumblebee | Hymenoptera | SAMEA7520483 | Female |
| *Bombus hypnorum* | Tree bumblebee | Hymenoptera | SAMEA7520655 | Male |
| *Bombus pascuorum* | Common carder bee | Hymenoptera | SAMEA7520484 | Female |
| *Bombus pratorum* | Early bumblebee | Hymenoptera | SAMEA7520485 | Female |
| *Bombus sylvestris* | Forest cuckoo bee | Hymenoptera | SAMEA7520657 | Male |
| *Bombus terrestris* | buff-tailed bumblebee | Hymenoptera | SAMEA7520487 | Female |
| *Nomada fabriciana* | Fabricius' Nomad Bee | Hymenoptera | SAMEA7520701 | Female |
| *Hylaeus communis* | Common Yellow-face Bee | Hymenoptera | SAMEA7746754 | Female |
| *Cerceris rybyensis* | Ornate Tailed Digger Wasp | Hymenoptera | SAMEA7701329 | Female |
| *Ectemnius continuus* | | Hymenoptera | SAMEA7520490 | Female |
| *Ectemnius lituratus* | | Hymenoptera | SAMEA7520491 | Female |
| *Mimumesa dahlbomi* | | Hymenoptera | SAMEA8603165 | Male |
| *Nysson spinosus* | Large Spurred Digger Wasp | Hymenoptera | SAMEA7520702 | Female |
| *Pemphredon lugubris* | Mournful Wasp | Hymenoptera | SAMEA8603139 | Unknown |
| *Trypoxylon clavicerum* | Club Horned Wood Borer Wasp | Hymenoptera | SAMEA7701565 | Female |
| *Myrmica sabuleti* | | Hymenoptera | SAMEA7520497 | Female |
| *Lasioglossum calceatum* | Common furrow bee | Hymenoptera | SAMEA7849393 | Unknown |
| *Lasioglossum lativentre* | Furry-claspered furrow bee | Hymenoptera | SAMEA7746765 | Male |
| *Lasioglossum leucozonium* | White-zoned furrow bee | Hymenoptera | SAMEA7746759 | Male |
| *Lasioglossum malachurum* | Sharp-collared furrow bee | Hymenoptera | SAMEA7746751 | Female |
| *Lasioglossum morio* | Common green furrow bee | Hymenoptera | SAMEA7746456 | Male |
| *Lasioglossum pauxillum* | Base-banded furrow bee | Hymenoptera | SAMEA7520494 | Female |

| | | | | |
|---|---|---|---|---|
| *Seladonia tumulorum* | Bronze furrow bee | Hymenoptera | SAMEA7746445 | Male |
| *Sphecodes ephippius* | Bare-saddled blood bee | Hymenoptera | SAMEA7746758 | Male |
| *Sphecodes monilicornis* | Box-headed blood bee | Hymenoptera | SAMEA7746755 | Male |
| *Amblyteles armatorius* | | Hymenoptera | SAMEA7520946 | Male |
| *Buathra laborator* | | Hymenoptera | SAMEA8534297 | Unknown |
| *Ichneumon xanthorius* | | Hymenoptera | SAMEA7746465 | Female |
| *Ophion luteus* | | Hymenoptera | SAMEA8534284 | Female |
| *Scambus nigricans* | | Hymenoptera | SAMEA7849231 | Female |
| *Megachile ligniseca* | Wood-carving leaf-cutter bee | Hymenoptera | SAMEA7520495 | Female |
| *Megachile willughbiella* | Willughby's leaf-cutter bee | Hymenoptera | SAMEA7520496 | Female |
| *Macropis europaea* | Yellow loosestrife Bee | Hymenoptera | SAMEA7746440 | Male |
| *Anoplius nigerrimus* | | Hymenoptera | SAMEA7746764 | Unknown |
| *Evagetes crassicornis* | | Hymenoptera | SAMEA7746601 | Unknown |
| *Athalia circularis* | | Hymenoptera | SAMEA7746777 | Unknown |
| *Athalia cordata* | | Hymenoptera | SAMEA7746763 | Unknown |
| *Athalia rosae* | Coleseed sawfly | Hymenoptera | SAMEA7520481 | Unknown |
| *Tenthredo livida* | | Hymenoptera | SAMEA7520699 | Unknown |
| *Tenthredo notha* | | Hymenoptera | SAMEA7746761 | Unknown |
| *Tiphia femorata* | Beetle killing wasp | Hymenoptera | SAMEA7520499 | Female |
| *Ancistrocerus nigricornis* | Early mason-wasp | Hymenoptera | SAMEA7746762 | Female |
| *Dolichovespula media* | Median wasp | Hymenoptera | SAMEA7520488 | Female |
| *Dolichovespula saxonica* | Saxon wasp | Hymenoptera | SAMEA7520489 | Male |
| *Dolichovespula sylvestris* | Tree wasp | Hymenoptera | SAMEA7746475 | Male |
| *Vespa crabro* | European hornet | Hymenoptera | SAMEA7520500 | Female |
| *Vespula germanica* | German wasp | Hymenoptera | SAMEA7520501 | Female |
| *Vespula vulgaris* | Common wasp | Hymenoptera | SAMEA7520502 | Female |
| *Chrysoperla carnea* | Common green lacewing | Neuroptera | SAMEA7520372 | Female |
| *Ischnura elegans* | Blue-tailed damselfly | Odonata | SAMEA7521125 | Female |
| *Sympetrum striolatum* | Common darter | Odonata | SAMEA7520376 | Female |
| *Chorthippus brunneus* | Field grasshopper | Orthoptera | SAMEA7520377 | Unknown |

| | | | | |
|---|---|---|---|---|
| *Chorthippus parallelus* | Meadow grasshopper | Orthoptera | SAMEA7520378 | Unknown |
| *Teleogryllus oceanicus* | Black field cricket | Orthoptera | SAMEA8023467 | Female |
| *Meconema thalassinum* | Oak bush-cricket | Orthoptera | SAMEA7520379 | Male |
| *Leuctra nigra* | | Plecoptera | SAMEA7521360 | Male |
| *Nemoura dubitans* | | Plecoptera | SAMEA9065873 | Female |
| *Nemurella pictetii* | | Plecoptera | SAMEA7520996 | Male |
| *Glyphotaelius pellucidus* | | Trichoptera | SAMEA7520965 | Male |
| *Limnephilus lunatus* | | Trichoptera | SAMEA7521206 | Female |
| *Limnephilus marmoratus* | | Trichoptera | SAMEA7520990 | Male |
| *Limnephilus rhombicus* | | Trichoptera | SAMEA7849396 | Male |
| *Polycentropus irroratus* | | Trichoptera | SAMEA9065840 | Unknown |
