## Supplemental Table 2: Overview detected Wolbachia genomes for "An endosymbiont harvest: Phylogenomic analysis of *Wolbachia* genomes from the Darwin Tree of Life biodiversity genomics project"

**Table S2: Overview of all detected *Wolbachia* infections**

| Scientific name host | Taxonomic order host | Wolbachia supergroup | Circular / linear | Coverage | Number of contigs | Genome size (bp) | BUSCO completeness | Assembly method | Estimated number of Wolbachia genomes | Estimated number of Wolbachia genomes per host |
|---|---|---|---|---|---|---|---|---|---|---|
| *Agriphila straminella* | Lepidoptera | B | circular | 42 | 1 | 1856314 | C:99.4%[S:98.9%,D:0.5%],F:0.0%,M:0.6%,n:364 | hifiasm | 2 | 2 |
| *Agriphila tristella* | Lepidoptera | B | linear | 135 | 2 | 1312870 | C:99.8%[S:99.5%,D:0.3%],F:0.0%,M:0.2%,n:364 | flye | 9 | 9 |
| *Aporia crataegi* | Lepidoptera | B | linear | 303 | 1 | 1310109 | C:99.7%[S:99.7%,D:0.0%],F:0.0%,M:0.3%,n:364 | flye | 6 | 6 |
| *Apotomis betuletana* | Lepidoptera | B | linear | 28 | 1 | 1635810 | C:99.7%[S:98.9%,D:0.8%],F:0.0%,M:0.3%,n:364 | hifiasm | 2 | 2 |
| *Apotomis turbidana* | Lepidoptera | B | circular | 51 | 1 | 1525756 | C:99.4%[S:98.9%,D:0.5%],F:0.3%,M:0.3%,n:364 | hifiasm | 3 | 3 |
| *Archips podanus* | Lepidoptera | B | circular | 50 | 1 | 1806407 | C:99.4%[S:98.9%,D:0.5%],F:0.3%,M:0.3%,n:364 | hifiasm | 4 | 4 |
| *Aricia agestis* | Lepidoptera | B | linear | 115 | 2 | 1562496 | C:99.7%[S:98.9%,D:0.8%],F:0.0%,M:0.3%,n:364 | hifiasm | 4 | 4 |
| *Autographa gamma* | Lepidoptera | Not well assembled - unclear number of Wolbachia strains | | | | | | | | |
| | | A | circular | 78 | 1 | 1594318 | C:99.5%[S:99.2%,D:0.3%],F:0.0%,M:0.5%,n:364 | flye | 4 | 9 |
| *Calamotropha paludella* | Lepidoptera | A | circular | 85 | 1 | 1378301 | C:99.5%[S:99.2%,D:0.3%],F:0.0%,M:0.5%,n:364 | flye | 5 | |
| *Campaea margaritaria* | Lepidoptera | B | circular | 456 | 1 | 1507682 | C:99.7%[S:99.2%,D:0.5%],F:0.0%,M:0.3%,n:364 | flye | 19 | 19 |
| *Camptogramma bilineatum* | Lepidoptera | B | circular | 41 | 1 | 1668206 | C:99.2%[S:98.4%,D:0.8%],F:0.0%,M:0.8%,n:364 | hifiasm | 2 | 2 |
| *Carcina quercana* | Lepidoptera | B | linear | 18 | 4 | 1572160 | C:99.8%[S:99.5%,D:0.3%],F:0.0%,M:0.2%,n:364 | flye | 1 | 1 |
| *Catoptria pinella* | Lepidoptera | B | circular | 83 | 1 | 1667357 | C:99.8%[S:99.5%,D:0.3%],F:0.0%,M:0.2%,n:364 | hifiasm | 3 | 3 |
| *Celastrina argiolus* | Lepidoptera | B | circular | 140 | 1 | 1329046 | C:99.7%[S:99.7%,D:0.0%],F:0.0%,M:0.3%,n:364 | hifiasm | 9 | 9 |
| *Chrysoteuchia culmella* | Lepidoptera | Not well assembled - unclear number of Wolbachia strains | | | | | | | | |
| *Colias croceus* | Lepidoptera | B | circular | 158 | 1 | 1396841 | C:99.8%[S:99.5%,D:0.3%],F:0.0%,M:0.2%,n:364 | hifiasm | 5 | 5 |
| *Cydia splendana* | Lepidoptera | A | linear | 154 | 2 | 1446399 | C:98.9%[S:98.6%,D:0.3%],F:0.3%,M:0.8%,n:364 | flye | 9 | 9 |
| *Emmelina monodactyla* | Lepidoptera | B | circular | 177 | 1 | 1385743 | C:99.7%[S:99.7%,D:0.0%],F:0.0%,M:0.3%,n:364 | hifiasm | 9 | 9 |
| | | A | circular | 18 | 1 | 1691857 | C:99.5%[S:99.2%,D:0.3%],F:0.0%,M:0.5%,n:364 | hifiasm | 2 | 4 |
| *Endotricha flammealis* | Lepidoptera | B | circular | 18 | 1 | 1747616 | C:99.7%[S:99.2%,D:0.5%],F:0.0%,M:0.3%,n:364 | hifiasm | 2 | |
| | | A | circular | 315 | 1 | 1431999 | C:99.2%[S:98.9%,D:0.3%],F:0.3%,M:0.5%,n:364 | hifiasm | 13 | 19 |
| *Epagoge grotiana* | Lepidoptera | A | circular | 135 | 1 | 1781589 | C:98.6%[S:98.6%,D:0.0%],F:0.5%,M:0.9%,n:364 | hifiasm | 6 | |
| *Epirrhoe alternata* | Lepidoptera | A | circular | 608 | 1 | 1197406 | C:98.9%[S:98.9%,D:0.0%],F:0.0%,M:1.1%,n:364 | hifiasm | 23 | 23 |
| *Erebia ligea* | Lepidoptera | B | linear | 33 | 4 | 1520330 | C:98.9%[S:98.4%,D:0.5%],F:0.0%,M:1.1%,n:364 | flye | 2 | 2 |
| *Erynnis tages* | Lepidoptera | B | linear | 113 | 1 | 1532960 | C:99.8%[S:99.5%,D:0.3%],F:0.0%,M:0.2%,n:364 | hifiasm | 8 | 8 |
| *Eucosma cana* | Lepidoptera | B | circular | 133 | 1 | 1407592 | C:98.9%[S:98.4%,D:0.5%],F:0.5%,M:0.6%,n:364 | hifiasm | 6 | 6 |
| *Euphydryas aurinia* | Lepidoptera | B | circular | 209 | 1 | 1802478 | C:99.7%[S:98.6%,D:1.1%],F:0.0%,M:0.3%,n:364 | hifiasm | 12 | 12 |
| *Eupithecia tripunctaria* | Lepidoptera | A | circular | 31 | 1 | 1478720 | C:99.4%[S:98.9%,D:0.5%],F:0.0%,M:0.6%,n:364 | hifiasm | 2 | 2 |
| *Hamearis lucina* | Lepidoptera | Not well assembled - unclear number of Wolbachia strains | | | | | | | | |
| *Hedya salicella* | Lepidoptera | Not well assembled - unclear number of Wolbachia strains + not enough coverage | | | | | | | | |
| *Hesperia comma* | Lepidoptera | Not well assembled - not enough coverage | | | | | | | | |
| *Hylaea fasciaria* | Lepidoptera | B | circular | 772 | 1 | 1307822 | C:99.7%[S:99.7%,D:0.0%],F:0.0%,M:0.3%,n:364 | flye | 22 | 22 |
| *Hypena proboscidalis* | Lepidoptera | Not well assembled - high number of detected SNPs with 10x data | | | | | | | | |

| | | | | | | | | | | |
|---|---|---|---|---|---|---|---|---|---|---|
| *Idaea aversata* | Lepidoptera | B | linear | 135 | 2 | 1828659 | C:99.4%[S:98.9%,D:0.5%],F:0.0%,M:0.6%,n:364 | hifiasm | 5 | 5 |
| *Laothoe populi* | Lepidoptera | Not well assembled - unclear number of Wolbachia strains | | | | | | | | |
| *Leptidea sinapis* | Lepidoptera | B | linear | 77 | 4 | 1438040 | C:99.5%[S:99.2%,D:0.3%],F:0.3%,M:0.2%,n:364 | flye | 4 | 4 |
| *Lycaena phlaeas* | Lepidoptera | B | circular | 24 | 1 | 1312316 | C:99.7%[S:99.7%,D:0.0%],F:0.0%,M:0.3%,n:364 | hifiasm-meta | 1 | 1 |
| *Melanargia galathea* | Lepidoptera | B | circular | 94 | 1 | 1550695 | C:99.7%[S:99.2%,D:0.5%],F:0.0%,M:0.3%,n:364 | flye | 7 | 7 |
| | | B | circular | 161 | 1 | 1381694 | C:99.7%[S:99.7%,D:0.0%],F:0.0%,M:0.3%,n:364 | hifiasm | 8 | 10 |
| *Nymphalis c-album* | Lepidoptera | B | circular | 43 | 1 | 1236246 | C:99.7%[S:99.7%,D:0.0%],F:0.0%,M:0.3%,n:364 | hifiasm | 2 | |
| *Ochlodes sylvanus* | Lepidoptera | B | linear | 27 | 2 | 1713863 | C:99.4%[S:98.9%,D:0.5%],F:0.0%,M:0.6%,n:364 | flye | 2 | 2 |
| *Opisthograptis luteolata* | Lepidoptera | Not well assembled - unclear number of Wolbachia strains | | | | | | | | |
| *Orgyia antiqua* | Lepidoptera | B | linear | 258 | 1 | 1546986 | C:99.7%[S:99.2%,D:0.5%],F:0.0%,M:0.3%,n:364 | flye | 10 | 10 |
| *Pammene fasciana* | Lepidoptera | B | circular | 103 | 1 | 1266982 | C:99.7%[S:99.7%,D:0.0%],F:0.0%,M:0.3%,n:364 | hifiasm | 6 | 6 |
| *Pandemis cinnamomeana* | Lepidoptera | B | linear | 34 | 4 | 1779235 | C:99.7%[S:99.2%,D:0.5%],F:0.0%,M:0.3%,n:364 | flye | 2 | 2 |
| *Pandemis corylana* | Lepidoptera | B | circular | 193 | 1 | 1510264 | C:99.4%[S:98.6%,D:0.8%],F:0.3%,M:0.3%,n:364 | flye | 8 | 8 |
| *Parapoynx stratiotata* | Lepidoptera | B | circular | 14 | 1 | 1545403 | C:99.7%[S:99.2%,D:0.5%],F:0.0%,M:0.3%,n:364 | hifiasm | 1 | 1 |
| *Pararge aegeria* | Lepidoptera | B | circular | 107 | 1 | 1332905 | C:99.7%[S:99.7%,D:0.0%],F:0.0%,M:0.3%,n:364 | hifiasm | 5 | 5 |
| | | A | circular | 62 | 1 | 1271809 | C:99.2%[S:98.9%,D:0.3%],F:0.0%,M:0.8%,n:364 | hifiasm | 4 | 21 |
| | | B | linear | 272 | 1 | 1358027 | C:99.5%[S:99.5%,D:0.0%],F:0.3%,M:0.2%,n:364 | hifiasm | 16 | |
| *Phalera bucephala* | Lepidoptera | B | linear | 9 | 1 | 1462983 | C:99.2%[S:98.9%,D:0.3%],F:0.3%,M:0.5%,n:364 | hifiasm | 1 | |
| *Pheosia gnoma* | Lepidoptera | A | circular | 393 | 1 | 1353731 | C:99.5%[S:99.5%,D:0.0%],F:0.0%,M:0.5%,n:364 | hifiasm | 13 | 13 |
| *Pheosia tremula* | Lepidoptera | B | circular | 243 | 1 | 1268782 | C:99.2%[S:99.2%,D:0.0%],F:0.5%,M:0.3%,n:364 | hifiasm | 7 | 7 |
| *Polyommatus icarus* | Lepidoptera | B | linear | 333 | 1 | 1574132 | C:99.8%[S:99.5%,D:0.3%],F:0.0%,M:0.2%,n:364 | hifiasm | 19 | 19 |
| *Pyrgus malvae* | Lepidoptera | B | circular | 254 | 1 | 1544873 | C:99.1%[S:98.6%,D:0.5%],F:0.5%,M:0.4%,n:364 | flye | 7 | 7 |
| *Rhopobota naevana* | Lepidoptera | B | circular | 13 | 1 | 1771993 | C:99.5%[S:98.4%,D:1.1%],F:0.0%,M:0.5%,n:364 | hifiasm | 1 | 1 |
| *Thymelicus sylvestris* | Lepidoptera | B | linear | 960 | 1 | 1286255 | C:99.7%[S:99.7%,D:0.0%],F:0.0%,M:0.3%,n:364 | flye | 48 | 48 |
| *Watsonalla binaria* | Lepidoptera | B | circular | 218 | 1 | 1651336 | C:99.7%[S:99.2%,D:0.5%],F:0.0%,M:0.3%,n:364 | hifiasm | 6 | 6 |
| *Xestia c-nigrum* | Lepidoptera | B | circular | 106 | 1 | 1372378 | C:99.7%[S:99.7%,D:0.0%],F:0.0%,M:0.3%,n:364 | hifiasm | 6 | 6 |
| *Yponomeuta plumbellus* | Lepidoptera | A | linear | 181 | 1 | 1418562 | C:98.9%[S:98.9%,D:0.0%],F:0.5%,M:0.6%,n:364 | hifiasm | 12 | 12 |
| *Apoderus coryli* | Coleoptera | A | circular | 31 | 1 | 1535196 | C:99.8%[S:99.5%,D:0.3%],F:0.0%,M:0.2%,n:364 | flye | 1 | 1 |
| *Nebria salina* | Coleoptera | Not well assembled - very high coverage (>1000x) | | | | | | | | |
| *Rhinocyllus conicus* | Coleoptera | A | circular | 123 | 1 | 1703908 | C:99.2%[S:98.4%,D:0.8%],F:0.3%,M:0.5%,n:364 | flye | 5 | 5 |
| | | A | circular | 25 | 1 | 1811324 | C:99.4%[S:98.9%,D:0.5%],F:0.0%,M:0.6%,n:364 | hifiasm | 2 | 4 |
| *Philonthus cognatus* | Coleoptera | B | linear | 25 | 2 | 1680410 | C:99.8%[S:98.4%,D:1.4%],F:0.0%,M:0.2%,n:364 | hifiasm | 2 | |
| *Acrocera orbiculus* | Diptera | A | circular | 121 | 1 | 1383045 | C:98.6%[S:98.6%,D:0.0%],F:0.3%,M:1.1%,n:364 | hifiasm | 2 | 2 |
| *Bombylius major* | Diptera | A | circular | 173 | 1 | 1556587 | C:99.4%[S:98.6%,D:0.8%],F:0.0%,M:0.6%,n:364 | flye | 5 | 5 |
| | | A | circular | 117 | 1 | 1434286 | C:99.2%[S:99.2%,D:0.0%],F:0.3%,M:0.5%,n:364 | flye | 6 | 21 |
| *Protocalliphora azurea* | Diptera | B | circular | 291 | 1 | 1550351 | C:99.5%[S:99.2%,D:0.3%],F:0.3%,M:0.2%,n:364 | flye | 15 | |
| *Sicus ferrugineus* | Diptera | A | circular | 119 | 1 | 1273941 | C:99.5%[S:99.5%,D:0.0%],F:0.3%,M:0.2%,n:364 | hifiasm | 5 | 5 |
| *Thecophora atra* | Diptera | Not well assembled - unclear number of Wolbachia strains | | | | | | | | |
| *Coremacera marginata* | Diptera | A | circular | 115 | 1 | 1387347 | C:99.2%[S:99.2%,D:0.0%],F:0.3%,M:0.5%,n:364 | hifiasm | 9 | 9 |
| *Baccha elongata* | Diptera | Not well assembled - unclear number of Wolbachia strains | | | | | | | | |

| | | | | | | | | | | |
|---|---|---|---|---|---|---|---|---|---|---|
| | | A | circular | 86 | 1 | 1368131 | C:99.4%[S:98.9%,D:0.5%],F:0.0%,M:0.6%,n:364 | hifiasm | 4 | 9 |
| *Cheilosia soror* | Diptera | A | linear | 96 | 2 | 1493041 | C:99.2%[S:99.2%,D:0.0%],F:0.0%,M:0.8%,n:364 | hifiasm | 5 | |
| *Cheilosia vulpina* | Diptera | Not well assembled - unclear number of Wolbachia strains | | | | | | | | |
| *Chrysotoxum verralli* | Diptera | Not well assembled - unclear number of Wolbachia strains | | | | | | | | |
| | | A | circular | 23 | 1 | 1483878 | C:99.4%[S:98.6%,D:0.8%],F:0.3%,M:0.3%,n:364 | flye | 2 | 3 |
| *Epistrophe grossulariae* | Diptera | A | circular | 20 | 1 | 1367569 | C:99.2%[S:98.9%,D:0.3%],F:0.3%,M:0.5%,n:364 | flye | 1 | |
| *Episyrphus balteatus* | Diptera | B | circular | 80 | 1 | 1402289 | C:99.2%[S:98.4%,D:0.8%],F:0.5%,M:0.3%,n:364 | hifiasm | 6 | 6 |
| *Eupeodes latifasciatus* | Diptera | B | circular | 406 | 1 | 1274269 | C:99.7%[S:99.7%,D:0.0%],F:0.0%,M:0.3%,n:364 | flye | 30 | 30 |
| *Melanostoma mellinum* | Diptera | B | circular | 43 | 1 | 1875946 | C:99.2%[S:98.4%,D:0.8%],F:0.0%,M:0.8%,n:364 | flye | 3 | 3 |
| *Platycheirus albimanus* | Diptera | A | linear | 12 | 6 | 1174338 | C:98.9%[S:98.9%,D:0.0%],F:0.5%,M:0.6%,n:364 | flye | 0 | 1 |
| | | A | circular | 37 | 1 | 1354583 | C:99.1%[S:98.6%,D:0.5%],F:0.5%,M:0.4%,n:364 | flye | 2 | 23 |
| *Sphaerophoria taeniata* | Diptera | B | linear | 375 | 3 | 1627501 | C:99.4%[S:98.6%,D:0.8%],F:0.3%,M:0.3%,n:364 | flye | 21 | |
| | | A | circular | 44 | 1 | 1502845 | C:98.6%[S:97.0%,D:1.6%],F:0.5%,M:0.9%,n:364 | hifiasm | 3 | 6 |
| *Volucella inflata* | Diptera | A | linear | 42 | 2 | 1346192 | C:99.5%[S:99.2%,D:0.3%],F:0.0%,M:0.5%,n:364 | hifiasm | 3 | |
| *Gymnosoma rotundatum* | Diptera | A | circular | 367 | 1 | 1431527 | C:98.9%[S:98.4%,D:0.5%],F:0.3%,M:0.8%,n:364 | flye | 24 | 24 |
| *Anomoia purmunda* | Diptera | A | circular | 177 | 1 | 1485792 | C:99.7%[S:99.2%,D:0.5%],F:0.0%,M:0.3%,n:364 | flye | 10 | 10 |
| *Merzomyia westermanni* | Diptera | A | circular | 163 | 1 | 1295458 | C:99.5%[S:99.2%,D:0.3%],F:0.0%,M:0.5%,n:364 | hifiasm | 10 | 10 |
| *Bibio marci* | Diptera | A | circular | 91 | 1 | 1355352 | C:98.7%[S:98.4%,D:0.3%],F:0.3%,M:1.0%,n:364 | flye | 3 | 3 |
| *Himacerus mirmicoides* | Hemiptera | Not well assembled - unclear number of Wolbachia strains | | | | | | | | |
| *Icerya purchasi* | Hemiptera | A | circular | 88 | 1 | 1372034 | C:99.4%[S:98.9%,D:0.5%],F:0.0%,M:0.6%,n:364 | hifiasm | 6 | 6 |
| | | A | circular | 31 | 1 | 1366845 | C:99.2%[S:99.2%,D:0.0%],F:0.3%,M:0.5%,n:364 | hifiasm | 2 | 3 |
| *Andrena dorsata* | Hymenoptera | A | circular | 11 | 1 | 1533669 | C:99.7%[S:99.2%,D:0.5%],F:0.0%,M:0.3%,n:364 | hifiasm | 1 | |
| *Andrena haemorrhoa* | Hymenoptera | A | circular | 194 | 1 | 1494006 | C:98.9%[S:98.6%,D:0.3%],F:0.5%,M:0.6%,n:364 | hifiasm | 5 | 5 |
| *Andrena hattorfiana* | Hymenoptera | A | linear | 7 | 3 | 1409226 | C:95.6%[S:95.3%,D:0.3%],F:2.7%,M:1.7%,n:364 | flye | 0 | 1 |
| *Nomada fabriciana* | Hymenoptera | A | circular | 321 | 1 | 1438975 | C:99.5%[S:99.2%,D:0.3%],F:0.0%,M:0.5%,n:364 | flye | 7 | 7 |
| *Hylaeus communis* | Hymenoptera | A | circular | 1030 | 1 | 1526906 | C:99.1%[S:98.6%,D:0.5%],F:0.3%,M:0.6%,n:364 | flye | 31 | 31 |
| *Ectemnius continuus* | Hymenoptera | A | circular | 349 | 1 | 1299221 | C:99.5%[S:99.2%,D:0.3%],F:0.0%,M:0.5%,n:364 | flye | 8 | 8 |
| *Trypoxylon clavicerum* | Hymenoptera | A | circular | 55 | 1 | 1287268 | C:98.6%[S:98.1%,D:0.5%],F:0.3%,M:1.1%,n:364 | flye | 1 | 1 |
| *Myrmica sabuleti* | Hymenoptera | A | linear | 49 | 1 | 1403111 | C:99.2%[S:99.2%,D:0.0%],F:0.3%,M:0.5%,n:364 | hifiasm | 2 | 2 |
| *Lasioglossum calceatum* | Hymenoptera | A | circular | 10 | 1 | 1271041 | C:98.7%[S:98.4%,D:0.3%],F:0.5%,M:0.8%,n:364 | flye | 1 | 1 |
| *Lasioglossum lativentre* | Hymenoptera | A | linear | 69 | 1 | 1306351 | C:99.5%[S:99.2%,D:0.3%],F:0.0%,M:0.5%,n:364 | flye | 4 | 4 |
| *Lasioglossum leucozonium* | Hymenoptera | Not well assembled - high number of detected SNPs with 10x data | | | | | | | | |
| | | A | linear | 359 | 2 | 1380835 | C:98.9%[S:98.6%,D:0.3%],F:0.3%,M:0.8%,n:364 | flye | 21 | 28 |
| *Lasioglossum malachurum* | Hymenoptera | A | linear | 119 | 2 | 1363507 | C:99.5%[S:99.5%,D:0.0%],F:0.0%,M:0.5%,n:364 | flye | 7 | |
| | | A | circular | 219 | 1 | 1282688 | C:99.5%[S:99.2%,D:0.3%],F:0.0%,M:0.5%,n:364 | hifiasm | 13 | 22 |
| | | A | circular | 72 | 1 | 1270694 | C:99.2%[S:99.2%,D:0.0%],F:0.0%,M:0.8%,n:364 | hifiasm | 4 | |
| *Lasioglossum morio* | Hymenoptera | A | linear | 79 | 1 | 1284395 | C:99.5%[S:99.2%,D:0.3%],F:0.0%,M:0.5%,n:364 | hifiasm | 5 | |
| *Seladonia tumulorum* | Hymenoptera | Not well assembled - unclear number of Wolbachia strains | | | | | | | | |
| *Sphecodes ephippius* | Hymenoptera | A | linear | 320 | 2 | 1317045 | C:99.1%[S:98.6%,D:0.5%],F:0.0%,M:0.9%,n:364 | hifiasm | 15 | 15 |
| | | A | circular | 245 | 1 | 1359653 | C:98.9%[S:98.9%,D:0.0%],F:0.5%,M:0.6%,n:364 | flye | 14 | 20 |

| | | | | | | | | | | |
|---|---|---|---|---|---|---|---|---|---|---|
| *Sphecodes monilicornis* | Hymenoptera | A | circular | 107 | 1 | 1141923 | C:99.5%[S:99.5%,D:0.0%],F:0.0%,M:0.5%,n:364 | flye | 6 | |
| *Scambus nigricans* | Hymenoptera | A | circular | 148 | 1 | 1150903 | C:98.9%[S:98.9%,D:0.0%],F:0.3%,M:0.8%,n:364 | hifiasm | 9 | 9 |
| *Macropis europaea* | Hymenoptera | A | circular | 74 | 1 | 1439111 | C:99.7%[S:99.7%,D:0.0%],F:0.0%,M:0.3%,n:364 | hifiasm | 5 | 5 |
| *Anoplius nigerrimus* | Hymenoptera | A | circular | 13 | 1 | 1345812 | C:98.3%[S:97.8%,D:0.5%],F:0.8%,M:0.9%,n:364 | flye | 1 | 1 |
| *Athalia cordata* | Hymenoptera | B | circular | 2302 | 1 | 1345842 | C:99.5%[S:98.4%,D:1.1%],F:0.3%,M:0.2%,n:364 | flye | 47 | 47 |
| *Tiphia femorata* | Hymenoptera | A | circular | 824 | 1 | 1330166 | C:98.1%[S:97.8%,D:0.3%],F:0.3%,M:1.6%,n:364 | hifiasm | 22 | 22 |
| *Ancistrocerus nigricornis* | Hymenoptera | A | circular | 549 | 1 | 1265149 | C:99.5%[S:99.5%,D:0.0%],F:0.0%,M:0.5%,n:364 | flye | 16 | 16 |
| *Dolichovespula media* | Hymenoptera | B | linear | 479 | 3 | 1685384 | C:99.1%[S:98.6%,D:0.5%],F:0.3%,M:0.6%,n:364 | flye | 13 | 13 |
| *Ischnura elegans* | Odonata | B | circular | 33 | 1 | 2192296 | C:99.5%[S:98.4%,D:1.1%],F:0.0%,M:0.5%,n:364 | hifiasm | 3 | 3 |
| *Sympetrum striolatum* | Odonata | A | linear | 113 | 1 | 1516595 | C:99.1%[S:98.6%,D:0.5%],F:0.3%,M:0.6%,n:364 | hifiasm | 6 | 6 |
| *Chorthippus brunneus* | Orthoptera | B | linear | 558 | 4 | 2013544 | C:99.5%[S:99.2%,D:0.3%],F:0.0%,M:0.5%,n:364 | flye | 33 | 33 |
| *Chorthippus parallelus* | Orthoptera | B | circular | 260 | 1 | 1748465 | C:99.5%[S:98.4%,D:1.1%],F:0.0%,M:0.5%,n:364 | flye | 20 | 20 |
