## Supplemental Table 3: Wolbachia reference genomes for "An endosymbiont harvest: Phylogenomic analysis of *Wolbachia* genomes from the Darwin Tree of Life biodiversity genomics project"

**Table S3: *Wolbachia* reference genomes**

| Assembly identifier | Short name | Taxonomic order host | Wolbachia supergroup | Circular / linear | Numbers of scaffolds | Genome size (bp) | BUSCO completeness | Removed contigs |
|---|---|---|---|---|---|---|---|---|
| GCA_902636475.1 | R_Col_DiaVir_A | Coleopta | A | linear | 105 | 1269619 | C:95.4%[S:95.1%,D:0.3%],F:2.5%,M:2.1%,n:364 | CACPRN010000079.1 (Brevundimonas), CACPRN010000081.1 (Pythium), CACPRN010000092.1 (Stenotrophomonas), CACPRN010000095.1(Rhizobacter), CACPRN010000100.1 (Sphingopyxis), CACPRN010000101.1 (Singulisphaera), CACPRN010000102.1 (Chitinophaga), CACPRN010000103.1 (Bacillus), CACPRN010000105.1 (uncultured bacterium), CACPRN010000106.1 (Sphingobacterium), CACPRN010000107.1 (Bosea), CACPRN010000109.1 (Advenella), CACPRN010000111.1 (Devosia), CACPRN010000119.1 (Niabella) |
| GCF_017896245.1 | R_Dip_AedAeg_A | Diptera | A | circular | 1 | 1267786 | C:99.5%[S:99.2%,D:0.3%],F:0.0%,M:0.5%,n:364 | |
| GCF_018454455.1 | R_Dip_CerCap_A | Diptera | A | linear | 65 | 1239646 | C:99.4%[S:98.9%,D:0.5%],F:0.0%,M:0.6%,n:364 | |
| GCA_021609905.1 | R_Dip_DelRad_A | Diptera | A | circular | 1 | 1586584 | C:98.9%[S:98.1%,D:0.8%],F:0.3%,M:0.8%,n:364 | |
| GCF_008033215.1 | R_Dip_DroAna_A | Diptera | A | circular | 1 | 1401460 | C:99.2%[S:99.2%,D:0.0%],F:0.3%,M:0.5%,n:364 | |
| GCA_014129655.1 | R_Dip_DroAra_A | Diptera | A | linear | 10 | 1293235 | C:99.5%[S:99.5%,D:0.0%],F:0.0%,M:0.5%,n:364 | |
| GCF_017916175.1 | R_Dip_DroAur_A | Diptera | A | linear | 116 | 1319217 | C:99.2%[S:99.2%,D:0.0%],F:0.0%,M:0.8%,n:364 | |
| GCF_014129605.1 | R_Dip_DroBai_A | Diptera | A | linear | 26 | 1190958 | C:97.5%[S:97.5%,D:0.0%],F:0.3%,M:2.2%,n:364 | |
| GCF_014129645.1 | R_Dip_DroBic_A | Diptera | A | linear | 24 | 1182871 | C:99.2%[S:99.2%,D:0.0%],F:0.0%,M:0.8%,n:364 | |
| GCF_014129685.1 | R_Dip_DroBif_A | Diptera | A | linear | 17 | 1190237 | C:99.2%[S:99.2%,D:0.0%],F:0.0%,M:0.8%,n:364 | |
| GCF_014129615.1 | R_Dip_DroBor_A | Diptera | A | linear | 16 | 1214204 | C:98.4%[S:98.4%,D:0.0%],F:0.0%,M:1.6%,n:364 | |
| GCF_001758565.1 | R_Dip_DroInc_A | Diptera | A | circular | 1 | 1267840 | C:95.6%[S:95.6%,D:0.0%],F:2.5%,M:1.9%,n:364 | |
| GCF_016584355.1 | R_Dip_DroMel_A | Diptera | A | circular | 1 | 1330657 | C:99.5%[S:99.2%,D:0.3%],F:0.0%,M:0.5%,n:364 | |
| GCF_014129535.1 | R_Dip_DroNeo_A | Diptera | A | linear | 19 | 1355678 | C:99.2%[S:99.2%,D:0.0%],F:0.0%,M:0.8%,n:364 | |
| GCF_014107455.1 | R_Dip_DroNik_A | Diptera | A | circular | 1 | 1139744 | C:99.7%[S:99.7%,D:0.0%],F:0.0%,M:0.3%,n:364 | |
| GCF_014129565.1 | R_Dip_DroOri_A | Diptera | A | linear | 19 | 1363981 | C:99.2%[S:98.9%,D:0.3%],F:0.0%,M:0.8%,n:364 | |
| GCA_000742435.1 | R_Dip_DroRec_A | Diptera | A | linear | 43 | 1126656 | C:99.2%[S:98.9%,D:0.3%],F:0.3%,M:0.5%,n:364 | |
| GCF_018467135.1 | R_Dip_DroSan_A | Diptera | A | circular | 1 | 1409094 | C:99.5%[S:99.5%,D:0.0%],F:0.0%,M:0.5%,n:364 | |
| GCA_014354315.1 | R_Dip_DroSec_A | Diptera | A | linear | 83 | 1294885 | C:99.4%[S:98.9%,D:0.5%],F:0.0%,M:0.6%,n:364 | |
| GCF_000022285.1 | R_Dip_DroSim_A | Diptera | A | circular | 1 | 1445873 | C:98.9%[S:98.9%,D:0.0%],F:0.5%,M:0.6%,n:364 | |
| GCF_014107475.1 | R_Dip_DroStu_A | Diptera | A | circular | 1 | 1185354 | C:99.7%[S:99.2%,D:0.5%],F:0.0%,M:0.3%,n:364 | |
| GCA_002300525.1 | R_Dip_DroSub_A | Diptera | A | linear | 106 | 1420341 | C:99.2%[S:99.2%,D:0.0%],F:0.3%,M:0.5%,n:364 | |
| GCA_000333795.2 | R_Dip_DroSuz_A | Diptera | A | linear | 110 | 1415350 | C:99.2%[S:99.2%,D:0.0%],F:0.3%,M:0.5%,n:364 | |
| GCA_005862135.1 | R_Dip_DroTei_A | Diptera | A | linear | 122 | 1301330 | C:99.5%[S:99.5%,D:0.0%],F:0.0%,M:0.5%,n:364 | |
| GCF_014129515.1 | R_Dip_DroTri_A | Diptera | A | linear | 9 | 1289544 | C:98.9%[S:98.9%,D:0.0%],F:0.0%,M:1.1%,n:364 | |
| GCF_014129525.1 | R_Dip_DroTro_A | Diptera | A | linear | 13 | 1215752 | C:99.5%[S:99.5%,D:0.0%],F:0.0%,M:0.5%,n:364 | |

| | | | | | | | | |
|---|---|---|---|---|---|---|---|---|
| GCA_005862115.1 | R_Dip_DroYak_A | Diptera | A | linear | 118 | 1288853 | C:99.5%[S:99.5%,D:0.0%],F:0.0%,M:0.5%,n:364 | |
| GCA_009732755.1 | R_Dip_HaeIrr_A | Diptera | A | circular | 1 | 1352354 | C:99.5%[S:99.5%,D:0.0%],F:0.0%,M:0.5%,n:364 | |
| GCF_018454475.1 | R_Dip_RagCer1_A | Diptera | A | linear | 16 | 1255676 | C:99.5%[S:99.2%,D:0.3%],F:0.0%,M:0.5%,n:364 | |
| GCF_017604245.1 | R_Dip_RhaCin_A | Diptera | A | circular | 1 | 1538351 | C:99.5%[S:99.5%,D:0.0%],F:0.0%,M:0.5%,n:364 | |
| GCA_014333535.1 | R_Hym_AnoGra_A | Hymenoptera | A | linear | 96 | 1202684 | C:99.7%[S:99.7%,D:0.0%],F:0.0%,M:0.3%,n:364 | |
| GCF_010820705.1 | R_Hym_AptDen_A | Hymenoptera | A | linear | 49 | 1174448 | C:94.8%[S:92.6%,D:2.2%],F:1.4%,M:3.8%,n:364 | NZ_SDUV01000004.1, NZ_SDUV01000005.1, NZ_SDUV01000007.1, NZ_SDUV01000011.1, NZ_SDUV01000012.1, NZ_SDUV01000017.1, NZ_SDUV01000019.1, NZ_SDUV01000020.1, NZ_SDUV01000029.1, NZ_SDUV01000048.1, NZ_SDUV01000049.1 (mitochondrion host) |
| GCF_902713635.1 | R_Hym_CarObs_A | Hymenoptera | A | linear | 148 | 1194983 | C:99.2%[S:98.9%,D:0.3%],F:0.0%,M:0.8%,n:364 | |
| GCA_017869155.1 | R_Hym_CerSol_A | Hymenoptera | A | circular | 1 | 1206967 | C:98.9%[S:98.9%,D:0.0%],F:0.3%,M:0.8%,n:364 | |
| GCA_902648475.2 | R_Hym_DiaAll_A | Hymenoptera | A | linear | 169 | 1249264 | C:97.3%[S:97%,D:0.3%],F:1.4%,M:1.3%,n:364 | CACRJV020000057.1, CACRJV020000099.1, CACRJV020000109.1, CACRJV020000112.1, CACRJV020000115.1, CACRJV020000128.1, CACRJV020000140.1, CACRJV020000144.1, CACRJV020000153.1, CACRJV020000156.1 (Rickettsia) |
| GCA_902636375.1 | R_Hym_DufNov_A | Hymenoptera | A | linear | 221 | 1188140 | C:95.3%[S:95.3%,D:0.0%],F:3.0%,M:1.7%,n:364 | CACPRR010000135.1 (mitochondrion host) |
| GCA_902636315.1 | R_Hym_LasAlb_A | Hymenoptera | A | linear | 228 | 1195679 | C:97.6%[S:97.3%,D:0.3%],F:1.9%,M:0.5%,n:364 | CACPRV010000175.1 (Sodalis) |
| GCF_009012935.1 | R_Hym_NasOne_A | Hymenoptera | A | linear | 47 | 1293406 | C:95.6%[S:95.3%,D:0.3%],F:0.3%,M:4.1%,n:364 | |
| GCA_001983615.1 | R_Hym_NasVit_A | Hymenoptera | A | linear | 136 | 1198879 | C:99.5%[S:99.2%,D:0.3%],F:0.0%,M:0.5%,n:364 | MUJM01000137, MUJM01000138 (mitochondrion host), MUJM01000139-MUJM01000142 (Proteus) |
| GCA_001675785.1 | R_Hym_NomFer_A | Hymenoptera | A | linear | 231 | 1337641 | C:98.6%[S:98.1%,D:0.5%],F:0.5%,M:0.9%,n:364 | |
| GCF_001675695.1 | R_Hym_NomFla_A | Hymenoptera | A | linear | 167 | 1332780 | C:98.7%[S:98.4%,D:0.3%],F:0.3%,M:1.0%,n:364 | |
| GCA_001675715.1 | R_Hym_NomLeu_A | Hymenoptera | A | linear | 182 | 1366719 | C:98.9%[S:98.9%,D:0.0%],F:0.3%,M:0.8%,n:364 | |
| GCA_001675775.1 | R_Hym_NomPan_A | Hymenoptera | A | linear | 191 | 1344355 | C:98.6%[S:95.9%,D:2.7%],F:0.5%,M:0.9%,n:364 | |
| GCA_017869285.1 | R_Hym_WiePum_A | Hymenoptera | A | circular | 1 | 1278697 | C:99.4%[S:97.5%,D:1.9%],F:0.0%,M:0.6%,n:364 | |
| GCF_006542295.1 | R_Lep_CarSas_A | Lepidoptera | A | circular | 1 | 1449344 | C:99.2%[S:99.2%,D:0.0%],F:0.0%,M:0.8%,n:364 | |
| GCA_902646255.1 | R_Ara_TetUrt_B | Arachnida | B | linear | 149 | 1157560 | C:99.5%[S:99.5%,D:0.0%],F:0.0%,M:0.5%,n:364 | |
| GCF_020405475.1 | R_Col_TriCon_B | Coleopta | B | linear | 12 | 1418452 | C:96.9%[S:96.4%,D:0.5%],F:0.0%,M:3.1%,n:364 | |
| GCF_004171285.1 | R_Dip_AedAlb_B | Diptera | B | circular | 1 | 1484007 | C:99.2%[S:98.4%,D:0.8%],F:0.3%,M:0.5%,n:364 | |
| GCF_018491735.1 | R_Dip_AnoDem_B | Diptera | B | linear | 64 | 1232500 | C:98.9%[S:98.9%,D:0.0%],F:0.0%,M:1.1%,n:364 | |
| GCF_018491625.2 | R_Dip_AnoMou_B | Diptera | B | circular | 1 | 1121812 | C:98.9%[S:98.6%,D:0.3%],F:0.3%,M:0.8%,n:364 | |
| GCF_008245065.1 | R_Dip_ChrMeg_B | Diptera | B | circular | 1 | 1376868 | C:99.7%[S:98.9%,D:0.8%],F:0.0%,M:0.3%,n:364 | |
| GCA_000723225.2 | R_Dip_CulMol_B | Diptera | B | linear | 103 | 1435676 | C:97.8%[S:97.5%,D:0.3%],F:1.1%,M:1.1%,n:364 | |
| GCF_000073005.1 | R_Dip_CulQui_B | Diptera | B | circular | 1 | 1482455 | C:99.4%[S:98.9%,D:0.5%],F:0.3%,M:0.3%,n:364 | |
| GCF_004795975.1 | R_Dip_DroMau_B | Diptera | B | circular | 1 | 1273530 | C:99.2%[S:98.9%,D:0.3%],F:0.3%,M:0.5%,n:364 | |
| GCF_000376585.1 | R_Dip_DroSim_B | Diptera | B | circular | 1 | 1301823 | C:99.8%[S:99.5%,D:0.3%],F:0.0%,M:0.2%,n:364 | |
| GCF_018454445.1 | R_Dip_RagCer5_B | Diptera | B | linear | 57 | 1180723 | C:98.3%[S:97.8%,D:0.5%],F:0.5%,M:1.2%,n:364 | |

| | | | | | | | |
|---|---|---|---|---|---|---|---|
| GCF_003999585.1 | R_Hem_BemTab_B | Hemiptera | B | circular | 1 | 1306495 | C:96.4%[S:94.8%,D:1.6%],F:1.4%,M:2.2%,n:364 |
| GCF_013458815.1 | R_Hem_DiaCit_B | Hemiptera | B | circular | 1 | 1656288 | C:99.7%[S:99.2%,D:0.5%],F:0.0%,M:0.3%,n:364 |
| GCA_018224395.1 | R_Hem_HomVit_B | Hemiptera | B | linear | 1 | 1712771 | C:98.9%[S:97.5%,D:1.4%],F:0.3%,M:0.8%,n:364 |
| GCF_007115015.1 | R_Hem_LaoStr_B | Hemiptera | B | linear | 2 | 1786382 | C:99.4%[S:98.9%,D:0.5%],F:0.0%,M:0.6%,n:364 |
| GCF_007115045.1 | R_Hem_NilLug_B | Hemiptera | B | linear | 2 | 1542238 | C:99.5%[S:98.4%,D:1.1%],F:0.0%,M:0.5%,n:364 |
| GCF_006334525.1 | R_Hym_LepCla_B | Hymenoptera | B | linear | 46 | 1150755 | C:98.9%[S:98.4%,D:0.5%],F:0.3%,M:0.8%,n:364 |
| GCF_001439985.1 | R_Hym_TriPre_B | Hymenoptera | B | circular | 1 | 1133809 | C:99.5%[S:99.5%,D:0.0%],F:0.0%,M:0.5%,n:364 |
| GCF_001027565.1 | R_Iso_ArmVul_B | Isopoda | B | linear | 10 | 1663741 | C:97.8%[S:97.5%,D:0.3%],F:1.1%,M:1.1%,n:364 |
| GCF_020995475.1 | R_Lep_CorCep_B | Lepidoptera | B | circular | 1 | 1359904 | C:99.5%[S:99.5%,D:0.0%],F:0.0%,M:0.5%,n:364 |
| GCF_018555315.1 | R_Lep_EreCas_B | Lepidoptera | B | linear | 2 | 1423447 | C:94.5%[S:94.2%,D:0.3%],F:0.0%,M:5.5%,n:364 |
| GCF_000333775.1 | R_Lep_HypBol_B | Lepidoptera | B | linear | 144 | 1377933 | C:99.2%[S:98.9%,D:0.3%],F:0.3%,M:0.5%,n:364 |
| GCF_001266585.1 | R_Lep_OphBru_B | Lepidoptera | B | linear | 120 | 1121046 | C:98.1%[S:98.1%,D:0.0%],F:0.5%,M:1.4%,n:364 |
| GCA_902636855.1 | R_Lep_ParAeg_B | Lepidoptera | B | linear | 64 | 1268638 | C:99.5%[S:99.5%,D:0.0%],F:0.3%,M:0.2%,n:364 |
| GCA_002318985.1 | R_Lep_PluAus_B | Lepidoptera | B | linear | 95 | 1158805 | C:99.7%[S:99.7%,D:0.0%],F:0.0%,M:0.3%,n:364 |
| GCF_018141665.1 | R_Lep_SpoPic_B | Lepidoptera | B | circular | 1 | 1339720 | C:99.4%[S:98.9%,D:0.5%],F:0.0%,M:0.6%,n:364 |
| GCF_013365455.1 | R_Nem_DirImm_C | Nematoda | C | circular | 1 | 920122 | C:94.5%[S:94.5%,D:0.0%],F:0.3%,M:5.2%,n:364 |
| GCF_000306885.1 | R_Nem_OncOch_C | Nematoda | C | circular | 1 | 957990 | C:95.1%[S:95.1%,D:0.0%],F:0.0%,M:4.9%,n:364 |
| GCF_000530755.1 | R_Nem_OncVol_C | Nematoda | C | linear | 1 | 960618 | C:94.8%[S:94.8%,D:0.0%],F:0.3%,M:4.9%,n:364 |
| GCF_000008385.1 | R_Nem_BruMal_D | Nematoda | D | circular | 1 | 1080084 | C:98.6%[S:98.6%,D:0.0%],F:0.3%,M:1.1%,n:364 |
| GCF_012030695.1 | R_Nem_BruPah_D | Nematoda | D | circular | 1 | 1072967 | C:98.4%[S:98.4%,D:0.0%],F:0.3%,M:1.3%,n:364 |
| GCF_013366805.1 | R_Nem_LitBra_D | Nematoda | D | linear | 41 | 1046149 | C:94.5%[S:94.5%,D:0.0%],F:0.0%,M:5.5%,n:364 |
| GCF_013365435.1 | R_Nem_LitSig_D | Nematoda | D | circular | 1 | 1045802 | C:95.1%[S:95.1%,D:0.0%],F:0.3%,M:4.6%,n:364 |
| GCF_002204235.2 | R_Nem_WucBan_D | Nematoda | D | linear | 100 | 1060850 | C:95.3%[S:95.3%,D:0.0%],F:1.9%,M:2.8%,n:364 |
| GCA_019061405.1 | R_Ara_FraSet_E | Arachnida | E | linear | 30 | 1082514 | C:98.4%[S:98.1%,D:0.3%],F:0.3%,M:1.3%,n:364 |
| GCF_001931755.2 | R_Col_FolCan_E | Coleopta | E | circular | 1 | 1801626 | C:99.5%[S:99.2%,D:0.3%],F:0.0%,M:0.5%,n:364 |
| GCF_014534705.1 | R_Hem_PenNig_E | Hemiptera | E | linear | 182 | 1457187 | C:98.7%[S:98.4%,D:0.3%],F:0.8%,M:0.5%,n:364 |
| GCA_001752665.1 | R_Nem_PratPen_E | Nematoda | E | linear | 12 | 975127 | C:95.6%[S:95.1%,D:0.5%],F:2.7%,M:1.7%,n:364 |
| GCF_012277295.1 | R_Sip_CteFel_E | Siphonaptera | E | circular | 1 | 1495538 | C:98.9%[S:97.0%,D:1.9%],F:0.3%,M:0.8%,n:364 |
| GCF_000829315.1 | R_Hem_CimLex_F | Hemiptera | F | circular | 1 | 1250060 | C:97.8%[S:97.3%,D:0.5%],F:0.8%,M:1.4%,n:364 |
| GCA_902636395.1 | R_Hym_OsmCae_F | Hymenoptera | F | linear | 163 | 1232261 | C:98.7%[S:98.4%,D:0.3%],F:0.5%,M:0.8%,n:364 |
| GCF_013366855.1 | R_Nem_MadHie_F | Nematoda | F | linear | 208 | 1025329 | C:93.1%[S:92.6%,D:0.5%],F:3.8%,M:3.1%,n:364 |
| GCF_020278625.1 | R_Nem_ManOzz_F | Nematoda | F | linear | 93 | 1073310 | C:95.3%[S:95.3%,D:0.0%],F:1.6%,M:3.1%,n:364 |
| GCF_012277315.1 | R_Sip_CteFel_F | Siphonaptera | F | circular | 1 | 1201647 | C:99.5%[S:99.2%,D:0.3%],F:0.3%,M:0.2%,n:364 |
| GCF_013365475.1 | R_Nem_CruTub_J | Nematoda | J | circular | 1 | 863988 | C:94.0%[S:94.0%,D:0.0%],F:0.8%,M:5.2%,n:364 |
| GCF_013365495.1 | R_Nem_DipCau_J | Nematoda | J | circular | 1 | 863427 | C:92.3%[S:92.3%,D:0.0%],F:0.5%,M:7.2%,n:364 |
| GCA_013317055.1 | R_Ara_AtePolK5_S | Arachnida | S | linear | 200 | 1404177 | C:86.5%[S:83.2%,D:3.3%],F:5.8%,M:7.7%,n:364 |
| GCF_013309895.1 | R_Ara_AtePolK3_S | Arachnida | S | linear | 373 | 1445964 | C:88.5%[S:83.8%,D:4.7%],F:3.8%,M:7.7%,n:364 |
