## Supplemental Table 4: Prophage modules for "An endosymbiont harvest: Phylogenomic analysis of *Wolbachia* genomes from the Darwin Tree of Life biodiversity genomics project"

**Table S4: Gene modules used to identify prophage regions**

| Module | Gene family | Gene identifier |
|---|---|---|
| Connector /Baseplate | OG0000021 | gpI |
| Connector /Baseplate | OG0000032 | gpJ |
| Connector /Baseplate | OG0000034 | gpW |
| Connector /Baseplate | OG0000058 | PAAR |
| Connector /Baseplate | OG0000045 | gpV |
| Connector /Baseplate | OG0000048 | Collar |
| Connector /Baseplate | OG0000044 | gpZ |
| Connector /Baseplate | OG0000041 | gpFII |
| Head | OG0000036 | Major capsid protein GpE |
| Head | OG0000042 | Head decoration protein D |
| Head | OG0000035 | Minor capsid protein Orf7 |
| Head | OG0000025 | Phage portal protein, lambda family |
| Head | OG0000049 | Head-to-tail joining protein W |
| Head | OG0000030 | Phage terminase |
| Tail | OG0000068 | Tail sheath protein |
| Tail | OG0000064 | Major tail tube protein |
| Tail | OG0000075 | gpG/GT |
| Tail | OG0000059 | Phage tail tape measure protein |
| Tail | OG0000080 | gpX (Tail protein X) |
| Tail | OG0000066 | gpD (late control) |
| Fibre | OG0000088 | Hypothetical protein |
| Fibre | OG0000069 | Domain of unknown function DUF4815 |
| Fibre | OG0000077 | Hypothetical protein |
| Fibre | OG0000100 | Hypothetical protein |
| Fibre | OG0000079 | Hypothetical protein |
| Fibre | OG0000104 | Hypothetical protein |
